# Hibernating ribosomes tether to mitochondria as an adaptive response to cellular stress during glucose depletion

**DOI:** 10.1101/2023.10.08.561365

**Authors:** Olivier Gemin, Maciej Gluc, Michael Purdy, Higor Rosa, Moritz Niemann, Yelena Peskova, Simone Mattei, Ahmad Jomaa

**Author notes:** These authors contributed equally (Olivier Gemin, Maciej Gluc).

## Abstract

Cell survival under nutrient-deprived conditions relies on cells’ ability to adapt their organelles and to rewire their metabolic pathways. In the fission yeast *Schizosaccharomyces pombe*, nutrient depletion is an unfavorable condition for protein synthesis and triggers a response characterized by mitochondrial fragmentation and the sequestration of cytosolic ribosomes on mitochondria. The molecular mechanism underlying ribosomal sequestration remains elusive. In this study, we performed time-lapse *in situ* cryo-electron tomography and cryo-electron microscopy complemented by biochemical experiments to elucidate the molecular details of this adaptive response. Our analysis indicate that upon glucose depletion protein synthesis is halted, causing ribosomes to enter an inactive state characterized by a conformational change that obstructs the peptidyl transferase center. Our *in situ* experiments reveal the presence of oligomeric arrays of hibernating ribosomes tethered to the mitochondrial surface. Surprisingly, ribosomes bind to the outer mitochondrial membrane *via* the small ribosomal subunit, an interaction facilitated by the ribosomal protein RACK1-orthologue Cpc2. Our experiments show that ribosome tethering is important for cell survival under glucose depletion conditions. This study broadens our understanding of the cellular adaptations triggered by nutrient scarcity and the underlying molecular mechanisms that regulate cell quiescence.

---

Cells initiate adaptive responses to counteract cellular stress resulting from nutrient depletion. This is accomplished through a shift in metabolic pathways and through alterations in organellar dynamics (*1–4*). In certain instances, cells exit the division cycle and enter a quiescent state, enabling them to endure harsh environmental conditions (*5*, *6*). Once optimal growth conditions are restored, cells can reactivate cell division and energy-consuming metabolic pathways. The inability of cells to effectively respond to nutrient cues has been linked to human degenerative diseases, diabetes, and aging (*7–9*). Furthermore, adaptive responses to these conditions are frequently observed within tumor microenvironments and contribute to immune evasion (*10*).

In yeast, glucose starvation triggers cellular adaptation by limiting the movement of macromolecular assemblies (*6*, *11*). Under these conditions, protein synthesis becomes unfavorable at both the transcriptional and translational levels while proteasomal activity and autophagy are enhanced, leading to adaptive modifications in the endoplasmic reticulum and mitochondria (*4*, *12–15*). Mitochondrial fragmentation is a common response to nutrient depletion, changes in carbon source and metabolic shifts. It enables various processes such as localized signaling, fatty acid oxidation for ATP production and mitophagy for the removal of damaged mitochondria (*13*, *16*, *17*). In yeast, prolonged glucose depletion induces mitochondrial fragmentation (*13*, *18*) and the sequestration of cytosolic ribosomes on mitochondria (*18*). The underlying molecular mechanisms of this adaptive response and how it could promote cell survival during stress are not yet known. We set out to explain how cytosolic ribosomes bind fragmented mitochondria in response to cellular stress triggered by glucose depletion in fission yeast.

## Results

### Ribosomes isolated under glucose depletion adopt an inactive state characterized by conformational changes in the active site

In cells, the presence of polysomes consisting of multiple ribosomes concurrently translating a single mRNA provides a direct indication of ongoing protein synthesis. To investigate the status of protein synthesis in glucose-depleted cells, we initially conducted polysome profiling on cells grown at different initial glucose concentration. This method allowed quantifying the relative amounts of polysomes engaged in translation compared to 80S monosomes and ribosomal subunits (Fig. 1A). Our data shows the presence of polysomes in cells grown under nutrient-rich conditions in Yeast Extract with Supplements (YES) and in defined Edinburgh Minimal Medium (EMM) supplemented with 2% w/v glucose, indicating active protein synthesis at high glucose concentration (fig. S1). The level of protein synthesis was analyzed in cells grown in EMM under low glucose conditions (0.5% w/v glucose) over up to 7 days (Fig. 1B-C and fig S1). The polysome profiles of samples harvested on day 1 and day 3 of glucose depletion resembled those measured in high glucose conditions. However, a notable shift occurred starting from day 4, characterized by an accumulation of 80S monosomes and the concomitant reduction of polysomes abundance. By day 7, the polysome profiles of glucose-depleted cells predominantly exhibited 80S peaks and absence of polysomes implying the cessation of protein synthesis.

**Fig. 1.**
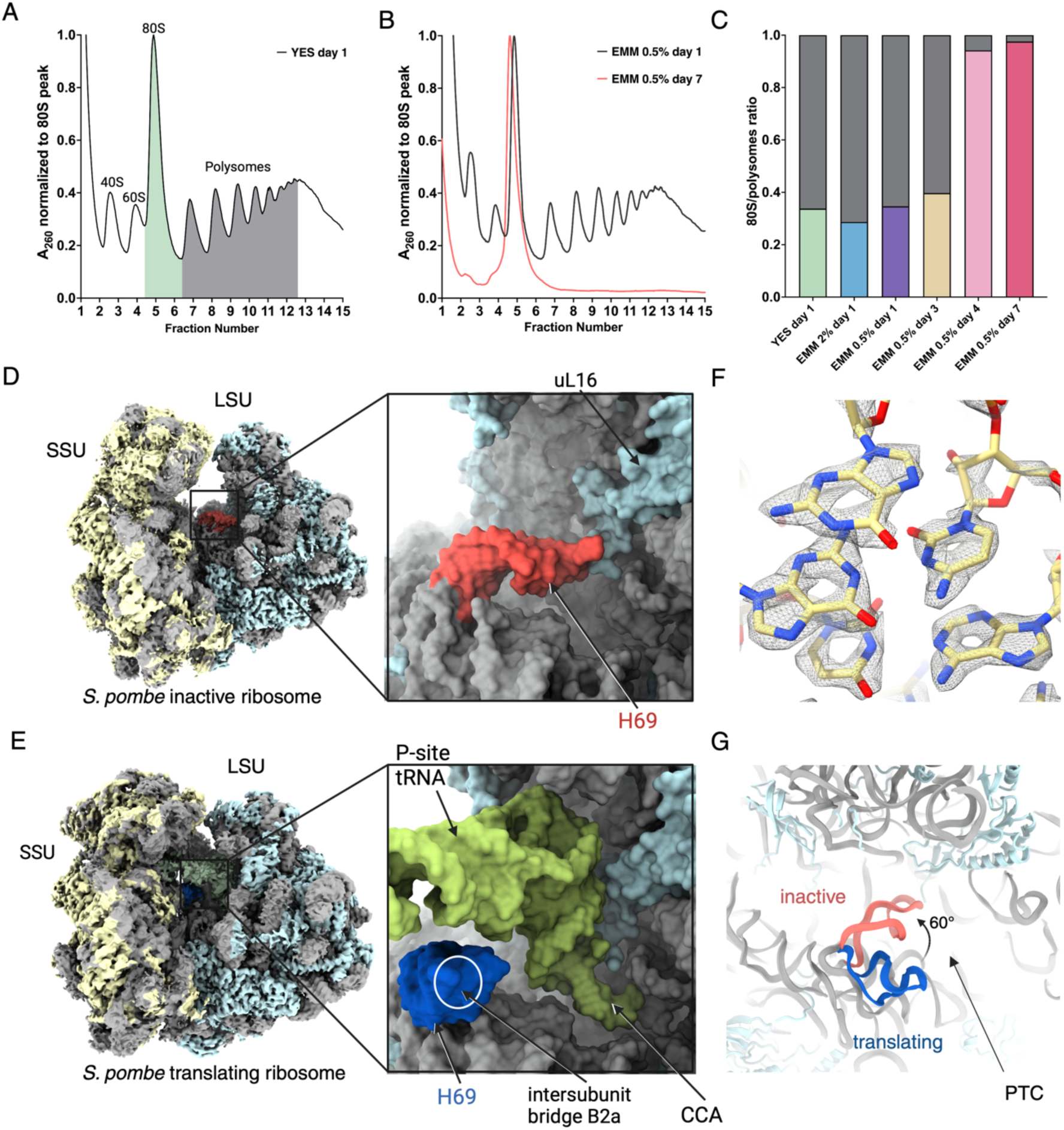
Biochemical and structural analysis of ribosomes isolated from cells cultured in glucose-depleted media reveals self-inhibition mode. (A) Profile of absorbance at 260 nm of a cell lysate obtained from an *S. pombe* culture grown in YES medium and then fractionated on a sucrose density gradient. Areas under the 80S monosome peak and the polysome peaks are highlighted green and gray, respectively. (B) Polysome profiles of *S. pombe* cells grown in EMM at a low glucose concentration and harvested at days 1 and 7 of glucose depletion. Signal normalized to the 80S peak absorbance. (C) Ratio of 80S to polysome quantities calculated from areas under the curves of the polysome profiles shown in (A), (B) and fig. S1. 80S fractions are shown for cells harvested after 1 day of growth in YES medium (green), in EMM with 2% w/v glucose (blue) and in EMM with 0.5% w/v glucose (purple). 80S fractions for glucose depleted cells after 3, 4 and 7 days of growth are shown in yellow, pink and red, respectively. The polysome fractions are shown in grey for all conditions. (D) Composite cryo-EM structure of the inactive *S. pombe* 80S ribosome isolated from cells at day 7 of glucose depletion. Inset shows a close-up of the peptidyl transferase center (PTC). H69 of the rRNA is highlighted in red. The SSU and LSU structures were separately refined using signal subtraction and focused 3D refinements, then displayed next to one another. Atomic coordinates are shown as surface. rRNA is colored gray and ribosomal proteins are colored cyan (LSU) and yellow (SSU). (E) Composite cryo-EM structure of the translating *S. pombe* 80S ribosome isolated from cells growing in glucose-rich media. Inset shows a close-up of the PTC. H69 of the rRNA is highlighted in blue. The SSU and LSU structures were separately refined using signal subtraction and focused 3D refinements, then displayed next to one another. Ribosomal proteins are colored cyan (LSU) and yellow (SSU), rRNA and tRNA are colored gray and green, respectively. (F) Close-up of the rRNA of the consensus 80S map refined to 1.9 Å resolution. EM-density is shown as mesh and atom coordinates are shown as sticks. (G) Close-up of the PTC depicting the conformational change in H69 observed between the translating ribosome (blue) and inhibited ribosome (red). The position of the polypeptide exit tunnel is indicated by an arrow.

To elucidate the mechanism of inhibition of cytosolic *S. pombe* ribosomes in glucose-depleted cells, we employed single-particle cryo-EM analysis (SPA) on biochemically isolated ribosomes. Initially, we determined a consensus reconstruction at a global resolution of 1.9 Å of crude cytosolic ribosomes isolated from cells cultured in EMM (0.5% glucose) at day 7 of glucose depletion (Fig. 1D, fig. S2 and table S1). For comparative purposes, we also determined a structure of the actively translating *S. pombe* ribosome isolated from cells growing in YES medium (Fig. 1E, fig. S3 and table S1). In the absence of deposited high-resolution structures of the *S. pombe* ribosome during the time of our study, we first built an atomic model of the *S. pombe* ribosome based on the obtained cryo-EM maps and homology modeling using the available structure of a cytosolic ribosome from *Saccharomyces cerevisiae* (Supplementary Material and Methods, Fig. 1F, fig. S4 and table S2). We built segments of the ribosome model *de novo*, revealing intriguing similarities to mammalian ribosomes (*19*). Notably, the ribosome protein L28e, absent in *S. cerevisiae* ribosomes (*20*), was detected in our structure and appeared to induce the ordering of the rRNA expansion segment ES7a upon binding (fig. S5). Moreover, the cryo-EM map revealed the presence of an additional 13 nucleotides that were not previously annotated as part of the 28S rRNA (fig. S6).

Ribosomes isolated from glucose-depleted cells were found to lack tRNA and mRNA, indicating a suppression of protein translation (Fig. 1D). Furthermore, our reconstruction revealed noticeable heterogeneity in both conformation and composition within the small ribosomal subunit (SSU) and disruptions in the peptidyl transferase center of the large subunit (LSU) (fig. S2). This heterogeneity was addressed using the data processing software cryoDRGN employing neural networks to parse the conformational heterogeneity of imaged ribosomes into representative 3D reconstructions (*21*) (fig. S7). Further 3D classification focused on the PTC and focused refinements improved the local resolution and revealed a conformational change in H69 in the PTC of the LSU (fig. S2). This helix exhibited a large rotation of 60**°** with respect to its position in the translating ribosome, bringing it closer to the extended loop of the ribosomal protein uL16 (Fig. 1D, E and G and fig. S8). Whereas in a translating ribosome these structural elements form a groove that facilitates LSU interaction with the CCA end of an incoming tRNA (*22*), here H69 occupies the P-site potentially preventing tRNA binding to the LSU due to steric clashes. Furthermore, the twisted configuration of H69 results in its displacement from its usual contact site with the SSU. This perturbation disrupts inter-subunit bridge B2a, a crucial element involved in initation of protein translation (*23*).

### Cytosolic hibernating ribosomes bind mitochondria via the small ribosomal subunit after prolonged glucose depletion

To elucidate the mechanism underlying how ribosomes become sequestered on the mitochondria in *S. pombe* under glucose depletion, we monitored changes at the ultrastructural and molecular level in the intracellular environment of wild-type (WT) cells cultured in defined EMM and subjected to glucose depletion over a period of 7 days. Daily measurement of the optical density of yeast cultures indicated that cells reached the stationary growth phase after 2-3 days (fig S9). Cell samples were collected at three distinct time points: day 0 (no glucose depletion) as well as day 4 and day 7 of glucose depletion. Cells were then vitrified on EM grids by plunge-freezing for subsequent structural analysis. Focused ion beam milling at cryogenic temperature (cryo-FIB)(*24*– *26*) was then used to obtain ≈200 nm-thick lamellae of the vitrified cells which were further imaged using cryo-electron tomography (cryo-ET) (Fig. 2A and B, figs. S9 and S10). The reconstructed tomograms exhibited mitochondrial fragmentation at day 4 but not at day 0, consistent with previous room-temperture electron tomography analyses of resin-embedded *S. pombe* samples (*18*). As glucose depletion progressed, mitochondria became entirely decorated with cytosolic ribosomes by day 7 (Fig. 2B and figs. S9 and S10).

**Fig. 2.**
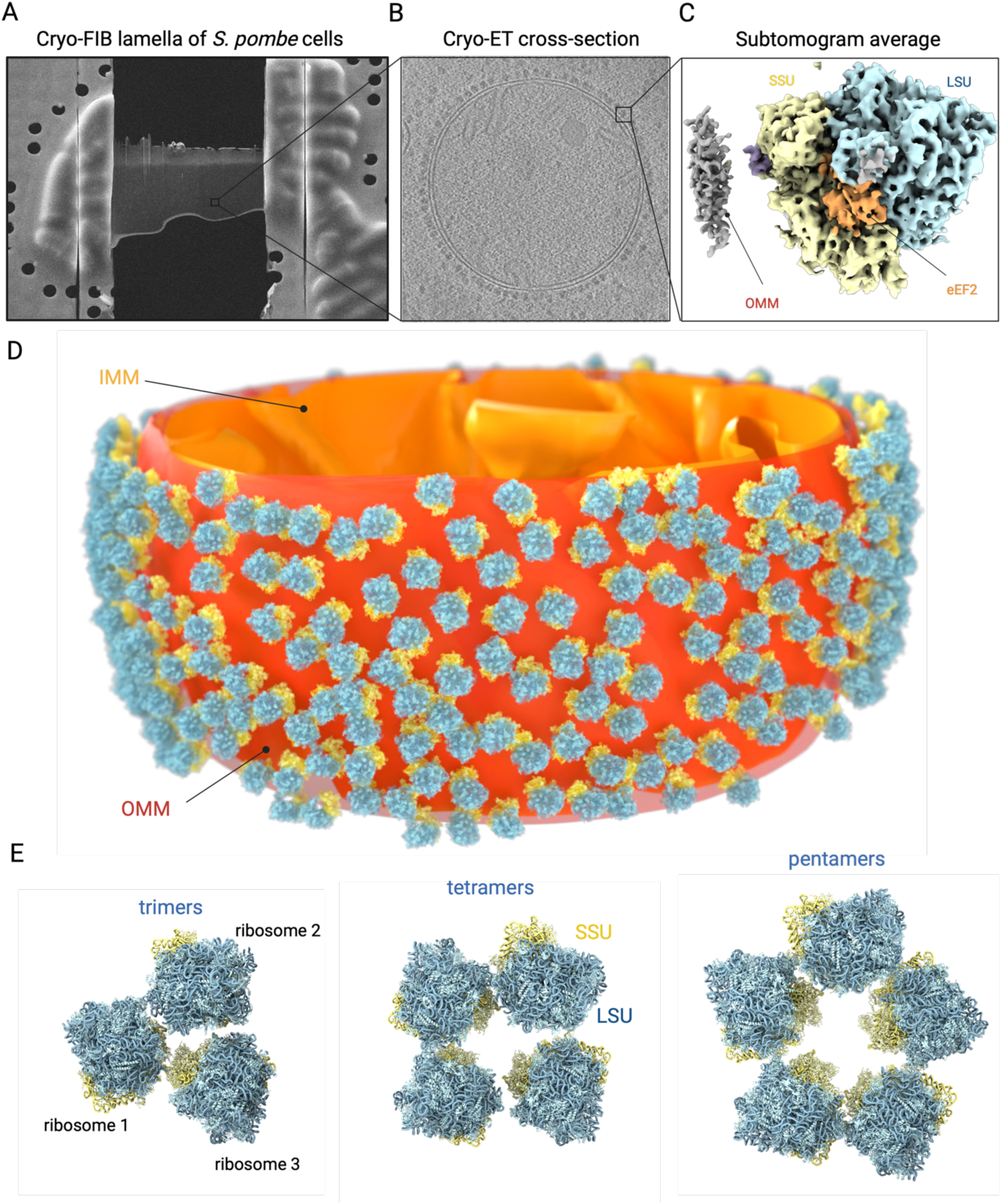
***In situ* cryo-ET and STA analysis of cytosolic hibernating ribosomes tethered to fragmented mitochondria after glucose depletion reveal oligomeric ribosome organization.** (A) Cultured *S. pombe* cells harvested at day 7 of glucose depletion, plunge-frozen on a cryo-EM grid and processed into a lamella using cryo-FIB-milling. Scale bar 20µm. (B) Computational slice through a representative tomographic reconstruction of a mitochondrion decorated with ribosomes in a *S. pombe* cell grown for 7 days in EMM at a low (0.5% w/v) glucose concentration. Scale bar 100nm. (C) Cryo-ET reconstruction of the hibernating ribosome at 11.4 Å resolution displayed as a surface and colored as yellow (SSU), cyan (LSU), purple (Cpc2) and orange (eEF2). (D) Segmentation of the mitochondrion shown in (B), superimposed with the position and orientation of OMM-bound ribosomes aligned by STA. The OMM is depicted in red and the IMM and its cristae are shown in orange. Ribosomal proteins and RNAs of the LSU and SSU are colored cyan and yellow, respectively. (E) Example arrays of hibernating ribosomes found on fragmented mitochondria, displayed as ribbons and color-coded as in (D).

We further processed our cryo-ET data by performing sub-tomogram averaging (STA)(*27*) on the subset of ribosomal particles that were docked at the outer mitochondrial membrane (OMM) (fig. S11). We aligned and averaged 10,192 OMM-associated ribosomes from 53 tomograms which resulted in an *in situ* cryo-ET structure of the tethered ribosome at 11.4 Å resolution (fig. S11 and table S3). The resulting density map showed that OMM-associated ribosomes are in a hibernating state, bound to eEF2, and devoid of tRNA (Fig. 2C and fig. S12). Mapping the coordinates of the aligned ribosomal particles back to their respective tomograms allowed analyzing their spatial distribution and orientation in the cellular context. Unexpectedly, the cytoplasmic ribosomes exclusively interacted with the OMM via their SSU (Fig. 2D and movie S1), in contrast to previous analyses of purified mitochondria from cycloheximide treated *S. cerevisiae*, where ribosomes were tethered *via* their large subunit with the ribosomal exit tunnel facing the OMM (*28*, *29*). The observed orientation is also distinct from translating ribosomes docked on the Sec translocon on the endoplasmic reticulum (ER) during cargo translocation across the ER membrane (fig. S13) (*30–34*). The map we obtained with STA does not show a clear connection between the 40S subunit and the OMM, suggesting that the tethering interaction is flexible and gets averaged out during sub-tomogram alignment.

Surprisingly, we observed an ordered organization of OMM-bound ribosomes in clusters of 2-to-5 ribosomes (Fig. 2D and E, and movie S1). These oligomeric assemblies revealed novel interactions between the neighboring ribosomes that had not been previously observed in either eukaryotic or bacterial ribosomal systems. The OMM-bound ribosome clusters were stabilized by interactions predominantly mediated by the ribosomal proteins eS7 and uS13 on the SSU, as well as eL27 and uL18 on the LSU (Fig. 3A and B). Notably, these protein pairs acted as pivotal hinge points, enabling the formation of the distinct oligomeric states observed on the OMM.

**Fig. 3.**
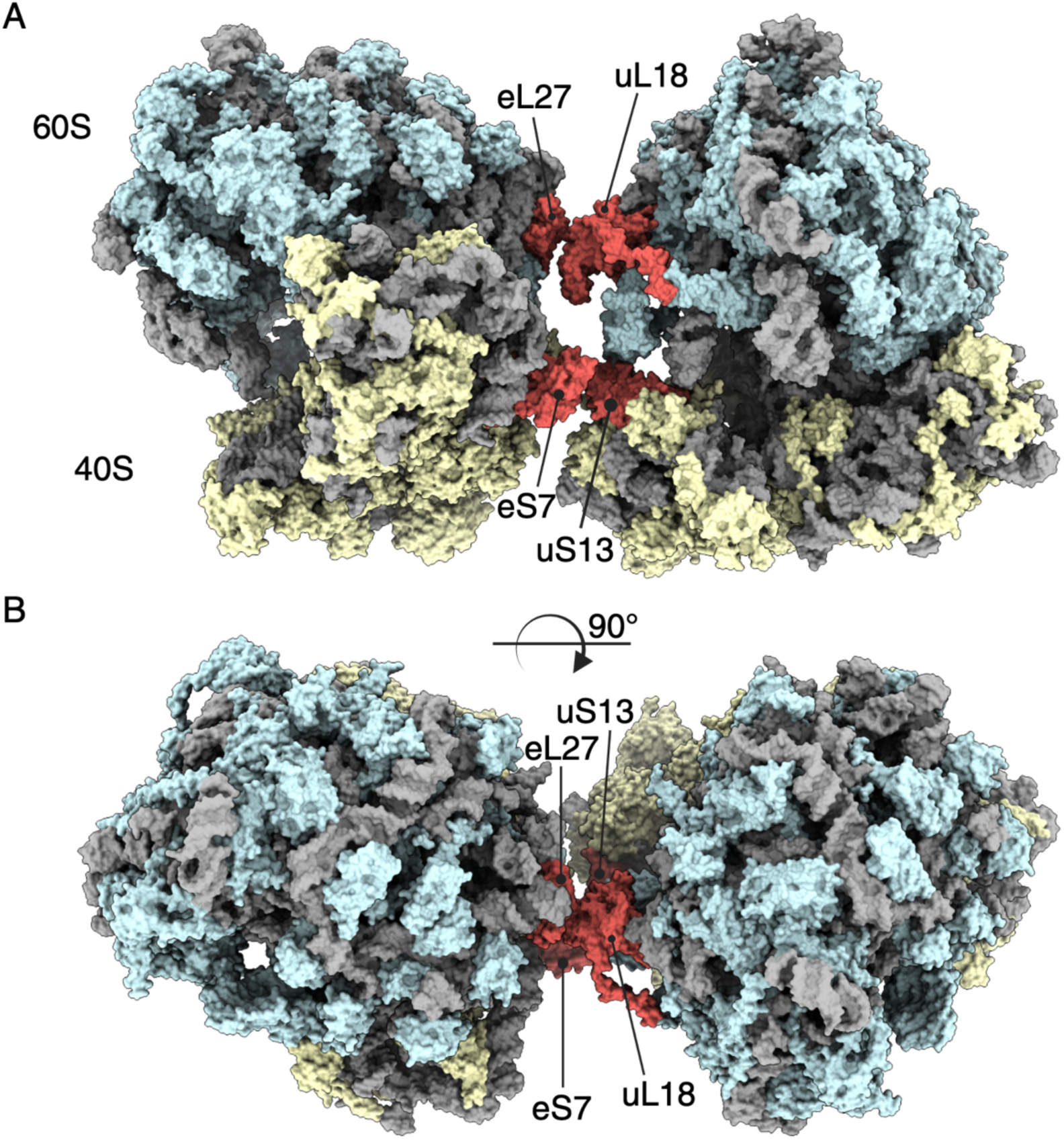
**Ribosome-ribosome interface in pentamers observed on the outer mitochondrial membrane.** (A) and (B) Interaction interface between hibernating ribosomes tethered to the OMM and engaged in a pentamer structure. The interactions are mediated via ribosomal proteins in a SSU-SSU and LSU-LSU interface. Models of the ribosomes are shown as surfaces, rRNA is colored gray, ribosomal proteins are colored cyan (LSU) and yellow (SSU) and the ribosomal proteins forming the interface (eL27, uL18, eS7, and uS13) are shown in red.

In summary, our results reveal that fragemented mitochondria become almost entirely decorated with hibernating ribosomes after prolonged glucose depletion of *S. pombe* cells. The hibernating ribosomes form oligomeric arrays that are tethered *via* the small ribosomal subunit to the OMM.

### Cpc2/RACK1 mediates the tethering of ribosomes to the OMM

To gain insights into the interaction between mitochondria and the hibernating ribosomes, we performed a rigid-body fit of the high-resolution structure of the cytosolic ribosome obtained using SPA cryo-EM into the cryo-ET map (Fig. 4A). Our analysis revealed that Cpc2 is present in close proximity to the OMM at an approximate distance of 30 Å. Cpc2 is known to transiently bind ribosomes where it is implicated in initiating protein synthesis, particularly for mitochondrial proteins (*35*, *36*). Cpc2 proximity to the OMM makes it an ideal candidate for mediating ribosome interactions with mitochondrial receptors to enhance cell persistence under glucose depletion.

**Fig. 4.**
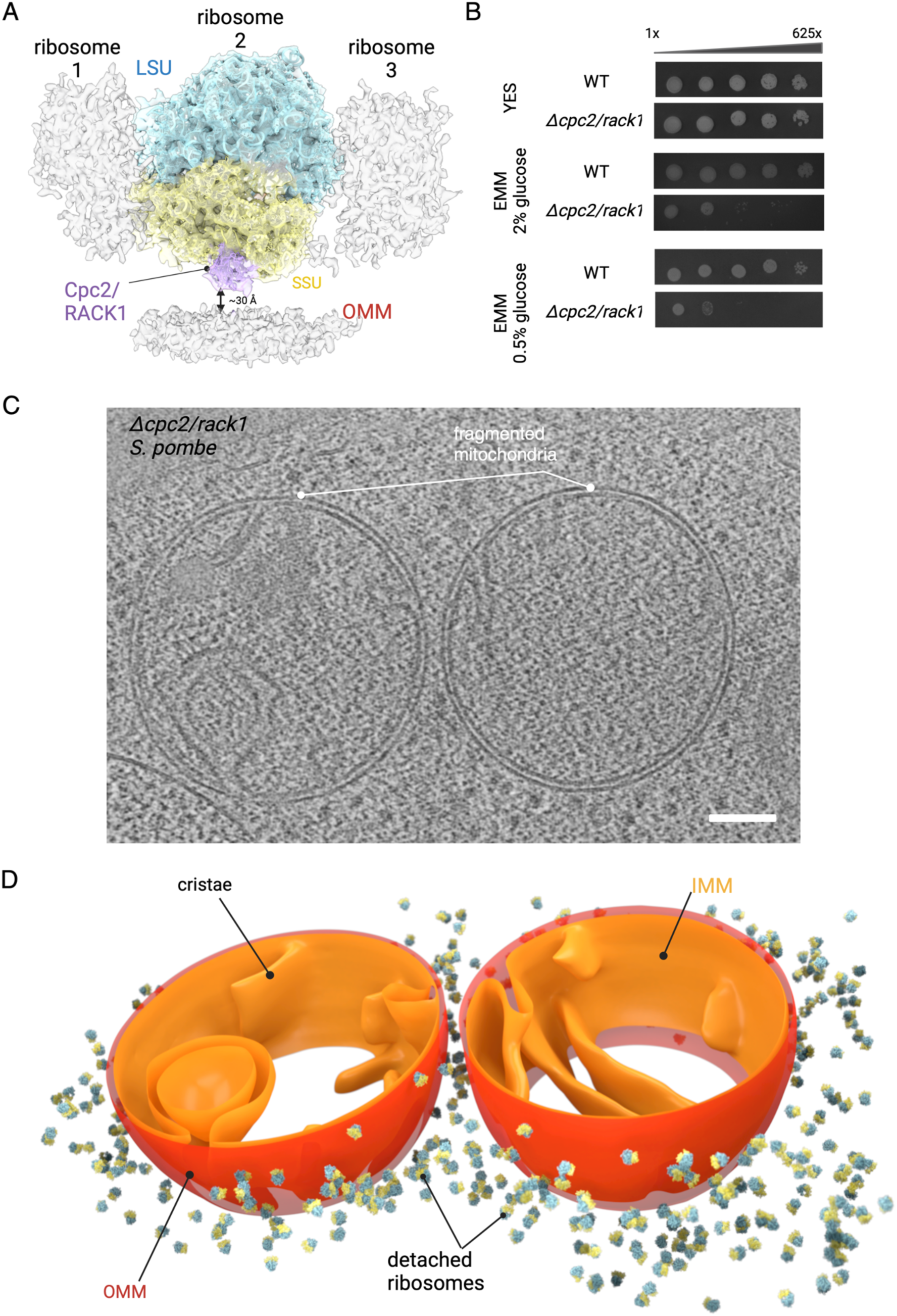
Cpc2/RACK1 regulates the tethering of cytosolic ribosomes to the OMM. (A) Cryo-EM structure of the *S. pombe* ribosome docked by rigid body fit in the cryo-ET reconstruction of a tethered ribosome at day 7 of glucose depletion, shown as a transparent surface. The arrow under the ribosome indicates the average distance separating RACK1 from the OMM. The position of two neighboring ribosomes (averaged out in the cryo-ET map) is indicated. (B) Viability assays of *S. pombe* WT and Δ*cpc2* strains cultured at 30 °C in either YES or EMM with high (2% w/v) or low (0.5% w/v) glucose concentration, and imaged after 3 days of growth. (C) Computational slice through a representative cryo-ET reconstruction of mitochondria in a Δ*cpc2* cell imaged at day 7 of glucose depletion; scale bar 100 nm. (D) Segmentation of the mitochondria shown in (C), superimposed with the position and orientation of cytosolic ribosomes aligned by STA. The OMM is depicted in red and the IMM and its cristae are shown in orange. Ribosomal proteins and RNAs of the LSU and SSU are colored cyan and yellow, respectively.

To validate our hypothesis, we investigated cell survival and ribosomal tethering in *S. pombe* strains lacking the ribosomal protein Cpc2 (Δ*cpc2*). We first monitored the survival of the Δ*cpc2* strain over several days using a dilution series viability assay (Fig. 4B). While Δ*cpc2* cells did not display discernible growth defects under nutrient-rich conditions, significant growth impairments were evident when these cells were cultivated in defined EMM media supplemented with either 2% or 0.5% glucose concentrations. We then used cryo-ET to assess whether ribosomal tethering to mitochondria was affected in the Δ*cpc2* strain. The tomograms showed fragmented circular mitochondria, as observed in WT cells, while no ribosome tethering was observed after 7 days of cells growing at low glucose concentrations (Fig. 4C-D, fig. S10 and movie S2).

Therefore, our data reveals a distinct mode of interaction between ribosomes and mitochondria, which is disrupted by the deletion of Cpc2. Our results indicate that Cpc2 is implicated in both cell viability and ribosome tethering to mitochondria under glucose depletion conditions, emphasizing its key role in mitochondrial homeostasis.

### Ribosome tethering is independent of mitochondrial fragmentation

Next, we investigated whether ribosome tethering was associated with mitochondrial fragmentation during glucose depletion. In *S. pombe*, fragmentation is mediated by the dynamin-related protein Dnm1, a homologue of the mammalian protein Drp1 (*13*). This GTPase accumulates on the OMM, triggers mitochondrial fission and is linked to mitochondrial dysfunction, fatty acid synthesis oxidation, mitophagy, apoptosis and glucose depletion (*37–40*).

We first evaluated cell viability during glucose depletion in a knockout strain of the protein Dnm1 (Δ*dnm1*). These cells did not display a growth defect when cultured in rich medium but they exhibited significantly slower growth rates in EMM at either glucose concentration (Fig. 5A). However, the growth defects were milder than those observed for the Δ*cpc2* strain. We then examined whether cytosolic ribosomes still tethered to mitochondria in Δ*dnm1* cells after prolonged glucose depletion. Cryo-ET imaging of Δ*dnm1* cells revealed that mitochondria remained elongated throughout glucose depletion (Fig. 5B). However, this morphological change did not prevent the tethering of cytosolic ribosomes to the OMM, with 98% of the imaged mitochondria (n=100) being fully decorated with ribosomes at day 7 of glucose depletion (Fig 5B, fig. S10 and movie S3).

**Fig. 5.**
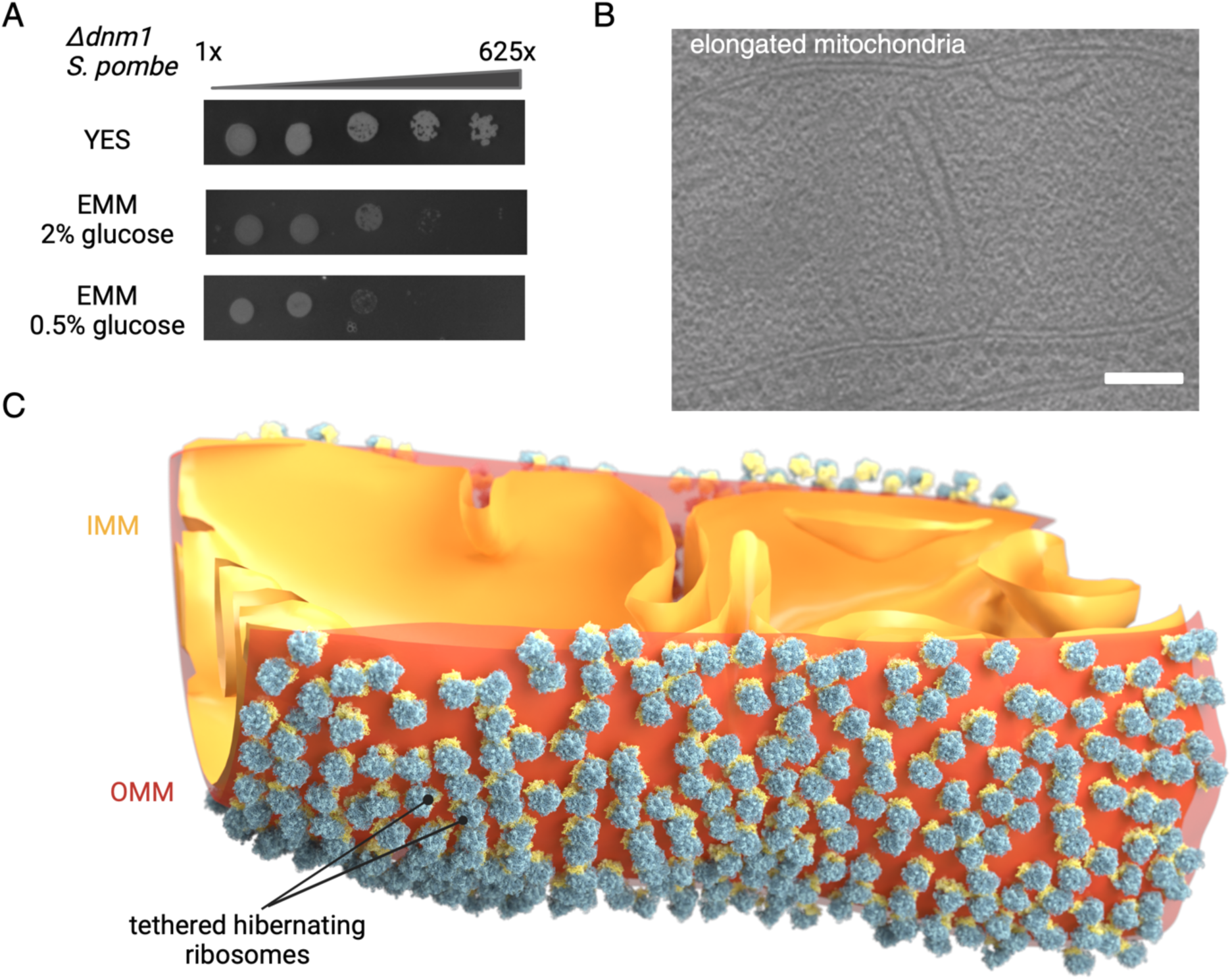
**Ribosome tethering is independent of Drp-1 mediated mitochondrial fragmentation.** (A) Viability assays of *S. pombe* WT and Δ*dnm1* strains cultured at 30 °C in either YES or EMM with high (2%) or low (0.5%) glucose concentration and imaged after 3 days of growth. (B) Computational slice through a representative cryo-ET reconstruction of a mitochondrion from a Δ*dnm1* cell imaged at day 7 of glucose depletion. Scale bar 100 nm. (C) Segmentation of the mitochondrion shown in (B), superimposed with the position and orientation of OMM-bound ribosomes aligned by STA. The OMM is depicted in red and the IMM and its cristae are shown in orange. Ribosomal proteins and RNAs of the LSU and SSU are colored cyan and yellow, respectively.

In conclusion, our results indicate that the process of ribosome tethering to the OMM in response to glucose starvation is independent of mitochondrial fragmentation, despite being subsequent to this morphological transition in wild type cells.

## Discussion

In this study, we describe a novel adaptive mechanism in yeast, by which arrays of hibernating cytosolic ribosomes become tethered to mitochondria in response to prolonged glucose depletion. Our biochemical analyses suggest a halt of protein synthesis by day 4 of glucose depletion, manifested by a decline in polysome amounts and an accumulation of 80S monosomes. Our cryo-EM structural analysis of cytosolic ribosomes isolated from glucose-depleted cells reveals an inactive ribosomal state lacking both tRNA and mRNA, in contrast to our structure of translating ribosomes isolated from cells grown in nutrient-rich media. The inactive ribosomes also exhibit conformational changes in the peptidyl transferase center, disrupting the inter-subunit bridge B2a which is integral for translation (*23*) and potentially preventing ribosomal interactions with either mRNA or tRNA. Such alterations in ribosome conformation likely mediate a downregulation of protein synthesis, enabling cells to sustain unfavorable conditions such as nutrient depletion.

Besides, our cryo-ET data showed notable morphological rearrangements including increased mitochondrial fragmentation and their reshaping into spheres, confirming previous observations of similar growth conditions (*13*, *18*). The recruitment of cytosolic ribosomes to the OMM occurred 6-7 days after glucose depletion and may have been triggered by an underlying signaling mechanism elicited 48-72 hours after the shutdown of protein synthesis (fig. S9). Cytosolic ribosomes tethered to the OMM self-assemble into distinct oligomeric arrays potentially interacting *via* ribosomal proteins eS7, uS13, eL27 and uL18 located on both ribosomal subunits. To our knowledge, this interaction interface has not been observed between eukaryotic ribosomes and it differs from interactions described between hibernating bacterial ribosomes (*41*), colliding ribosomes in yeast and mammals (fig. S14) (*42*) and ribosome dimers reported in mice and pathogens (*43*, *44*).

Our *in situ* structural analysis identifies Cpc2, an orthologue of the mammalian RACK1, as a key player in the tethering of ribosomes to the OMM *via* their SSU during glucose depletion. Cells lacking this protein fail to tether ribosomes to mitochondria and show reduced viability under depleted glucose conditions. RACK1/Cpc2 is an integral, eukaryotic-specific ribosomal protein required for the initiation of protein translation on short mRNAs including mitochondrial protein translation (*35*). In humans, RACK1 participates in regulating apoptosis by recruiting BAX and BIM to the OMM (*45*). In yeast, deletion of this protein results in the downregulation of mitochondrial proteins involved in respiration, oxidative stress response and iron homeostasis. It is thus possible that Cpc2 initially recruits ribosomes that are synthesizing mitochondrial proteins to the mitochondria (*29*, *34*), during the initial phase of switching from glycolysis to respiration (*46*). It is tempting to speculate that those ribosomes will then stay bound to the OMM via the Cpc2 interaction when translation is shut down due to glucose depletion.

In contrast, our cryo-ET data of Δ*dnm1* cells show that mitochondrial fission mediated by Dnm1 is uncoupled from the process of ribosome tethering. In eukaryotes, Dnm1/Drp1 is responsible for fission and it is upregulated in mitochondria undergoing fragmentation (*13*, *37–39*). Our biochemical experiments also indicate that cell viability is affected to a lesser extent in cells lacking this protein, which further corroborates that ribosome tethering plays a role in ensuring cell survival under glucose depletion.

Altogether, our biochemical and structural investigations reveal uncharacterized mitochondrial dynamics underlying a cellular response to nutrient scarcity, a condition reminiscent of disease models including cancer and diabetes, in which RACK1 and protein synthesis also play key roles (*47*). After prolonged nutrient depletion, mitochondria act as a cellular stress sensor and switch off their protein import to preserve energy and resources. We propose a model in which ribosomes stably bind mitochondria during cell quiescence and are then kept at the OMM in a hibernating state until nutrient conditions are restored. It is possible that tethered ribosomes perform a moonlighting function promoting cell survival by protecting fragmented mitochondria from autophagy and preventing apoptosis by stabilizing the mitochondrial membrane potential, thus averting cell death. Finally, since the effects of glucose starvation are reversible (*11*, *12*, *18*), we speculate that upon the reinstatement of high-nutrient conditions and the resurgence of protein synthesis, OMM-bound ribosomes would be ideally placed to synthesize mitochondrial proteins (*34*) to jumpstart mitochondrial activity. Upon the completion of one cycle of mitochondrial protein synthesis, OMM-bound ribosomes may then be released from mitochondria to translate new mRNA molecules.

In conclusion, our study uncovers a previously unexplored adaptive mechanism operating within a eukaryotic system to ensure persistence. This mechanism exemplifies energy regulation under environmental stress through the recruitment of the protein synthesis machinery at the mitochondria and the suspension of biosynthetic activity. This response is underpinned by rearrangements at the organellar and molecular levels and contributes to the initiation of cell dormancy.

Further investigations are needed to evaluate whether the recruitment of hibernating ribosomes to mitochondrial membranes is prompted by other environmental cues inducing cellular stress and if this mechanism is also conserved in mammalian systems. Since adaptive response to nutrients cues is commonly observed in cancer, our study could also provide the molecular understanding on cell persistance in pathologicial enviroments.

## Acknowledgments

We thank the Jomaa and Mattei lab members for discussions. We thank the Mahamid group members for fruitful discussions and help, especially Julia Mahamid and Ievgeniia Zagoriy for providing the *S. pombe* strain and for training on cryo-FIB milling, and Liang Xue for support with data analysis. We thank Kelly Dryden for support with cryo-EM data collection; Ricardo Sanchez and Amy Wang for support with data analysis. We thank Jochen Zimmer, Joaquin Ortega, Niccolò Banterle, Gautam Dey, and Julia Mahamid for critical reading of the manuscript. Single particle cryo-EM data collection was performed at the electron microscopy core facility MEMC at the School of Medicine, University of Virginia. We acknowledge the access and services provided by the Imaging Centre at the European Molecular Biology Laboratory (EMBL IC), generously supported by the Boehringer Ingelheim Foundation. We thank the EMBL central IT services and Thomas Hoffmann for computational support.

## Author contributions

A.J., S.M., O.G. and M.G. conceived the study, M.G. and M.N. purified the ribosomes and performed the biochemical experiments. O.G., M.G., M.N., and Y.P. performed yeast culture. O.G. and H.R. performed cryo-FIB milling and cryo-ET data collection and processing. M.G. and M.P. collected and processed the cryo-EM data. M.G. and A.J. built atomic models. A.J. and S.M. supervised the work. A.J., S.M., O.G., M.G. wrote the manuscript. All authors contributed to data analysis and the final version of the manuscript.

## Data and materials availability

Cryo-EM maps and model coordinates are deposited in the EMDB as EMD-XXXX, and EMD-XXXX, in the PDB as PDB ID XXX, XXX, for the *S. pombe* inactive and translating ribosomes, and EMDB-XXXX for the in-situ cryo-ET hibernating ribosome. All other data are available in the main text or the supplementary materials.

