## Supplementary Material for "Hibernating ribosomes tether to mitochondria as an adaptive response to cellular stress during glucose depletion"

† These authors contributed equally

### MATERIALS AND METHODS:

#### POLYSOME PROFILING ON SUCROSE GRADIENT

*Schizosaccharomyces pombe* cells were cultured at 30 °C under agitation in YES (Yeast Extract with Supplements) medium or in EMM (Edinburgh Minimal Medium) containing either 2% or 0.5% (w/v) glucose until they reached an optical density at 600 nm ($OD_{600}$) of 1, or in EMM media with 0.5% (w/v) glucose for durations of 3, 4, and 7 days. At the desired OD or time points, the cells were incubated with 100 µg/mL of cycloheximide for 15 minutes at 30 °C. Cells were then harvested and lysed using 0.5 mm glass beads. The absorbance at 260nm ($A_{260}$) of the cell lysate was measured after diluting the samples 50x with 0.1% SDS. All samples were then diluted to a concentration of 24 $A_{260}$ units per mL. 12 $A_{260}$ units of each sample were then loaded on a continuous 10-50% sucrose gradient and centrifuged at 230,000 g for 2.5 hours at 4 °C to separate ribosomes and poly-ribosomes. The poly-ribosome gradients were analyzed using a BIOCOMP Piston Gradient Fractionator™ system and visualized using the software GraphPad Prism 10.

#### RIBOSOME ISOLATION

An overnight culture of *S. pombe* cells was grown in 10 mL of either YES medium or EMM supplemented with 2% glucose (w/v). To obtain non-translating ribosomes, the cell culture was transferred to a culture flask containing 500 mL of EMM supplemented with 0.5% glucose (w/v), and then grown for 7 days. To obtain translating ribosomes, cells were transferred to 500 mL of YES medium and harvested on the following day. Cells were harvested by centrifugation at 3000 g for 5 minutes at room temperature and the resulting cell pellet was washed with Lysis Buffer (20mM Hepes pH 7.4, 100mM KCl, 5mM $Mg(OAc)_2$, protease inhibitor cocktail tablet) and transferred to a 50 mL centrifugation tube. The sample was spun at 5000 g for 3 minutes at 4 °C. The resulting pellets were resuspended in 4 mL of Lysis Buffer and transferred to a 15 mL centrifugation tube. The cells were then lysed using 0.5 mm glass beads by subjecting them to four cycles of vortexing at 4 °C for 30 seconds interspersed with 30 seconds of cooling on ice between each cycle. The cell lysate was centrifuged at 5000 g for 5 minutes at 4 °C and then clarified to remove cell debris by decanting. The resulting supernatant was transferred on a 50% sucrose cushion (50% w/v sucrose, 20mM Hepes pH 7.4, 60mM KCl, 5mM $Mg(OAc)_2$) in 13.5 mL quick-seal ultracentrifuge tubes. Subsequently, samples were centrifuged at 43,000 rpm (50.2 Ti rotor, 250,000 g) for 20 hours at 4 °C. Following centrifugation, ribosome pellets were carefully resuspended in 150 µL cryo-EM buffer (20mM Hepes pH 7.4, 60mM KCl, 5mM $Mg(OAc)_2$) and diluted to a concentration of 250 µg/µL as determined by spectrophotometric analysis. Freshly prepared ribosome samples were applied to cryoEM grids.

#### ELECTRON MICROSCOPY DATA COLLECTION

Five µL of either translating ribosomes isolated from rich media or non-translating ribosomes grown in EMM supplemented with 0.5% glucose (w/v) were applied to Quantifoil Cu200 R2/1 cryo-EM grids previously coated with a 2 nm layer of carbon using a Safematic carbon coater (Rave Scientific). Grids were first rendered hydrophilic by plasma cleaning at 15 mA for 15 s using Pelco easiGlow™ system before sample application. Sample was incubated on grids for 60 s and then blotted for 10 s with blot force +7 before plunge-freezing in liquid ethane using a

Vitrobot Mark IV (ThermoScientific) with the environmental chamber set to 4 °C and 100% humidity.

Data acquisition was carried out using a Titan Krios (ThermoScientific) transmission electron microscope (TEM) operated at an acceleration voltage of 300 kV and equipped with a K3 direct electron detector and a Quantum energy filter (Gatan). A total of 3,582 and 9,424 movie frames were acquired for translating and non-translating ribosome samples respectively, at a nominal magnification of 105,000x resulting in a calibrated pixel size of 0.83 Å. The TEM was operated with a spot size of 5, and each movie frame was exposed for 4.06 s, accumulating a total dose of 50 $e^{-}/Å^{2}$ in counting mode, using a filter slit size of 10 eV. The target defocus range was set between -1.6 and -0.6 µm. The acquired movie frames were motion-corrected and dose-weighted using the software cryoSPARC (*48*). The contrast transfer function (CTF) of dose-weighted micrographs was estimated using cryoSPARC.

**CRYO-EM DATA PROCESSING**

Cryo-EM data processing was carried out following the workflows depicted in figs. S2 and S3. CryoSPARC was employed for all processing steps. 3,582 micrographs of translating ribosomes were subjected to CTF estimation and automated particle picking to extract 315,521 initial particles. A subset of 226,425 ribosomal particles was selected using 2D classification. These particles underwent Heterogenous and Homogenous refinement in cryoSPARC, resulting in a consensus map at an average resolution of 2.13 Å. Following 3D variability analysis using a mask focused on the tRNA density, 58,737 particles exhibiting stable tRNA densities underwent a final round of Homogenous refinement, yielding a 80S ribosome map at an average resolution of 2.38 Å. Particle Subtraction masking either the 40S or the 60S ribosomal subunit granted a separate map of each subunit. The final 40S and 60S maps reached an average resolution of 2.84 Å and 2.41 Å respectively.

Following the CTF estimation of 9,424 micrographs of the non-translating ribosomes, automated particle picking allowed extracting 1,496,067 particles that were directly subjected to Heterogenous Refinement without prior 2D classification. From this dataset, the 820,453 particles showing the strongest SSU density underwent Homogenous and Non-uniform refinements, resulting in a consensus map at an average resolution of 1.94 Å. These particles were subjected to another round of Heterogenous Refinement to further improve the SSU density. Resulting classes exhibiting the strongest SSU density were selected, amounting to 293,749 particles. The structure reconstructed using those particles showed an unidentified density in the PTC of the LSU. To better resolve this density, particles were subjected to a 3D Variability analysis using a mask focused on the peptidyl transferase center (PTC). The particle class exhibiting H69 in a twisted conformation contained 73,376 particles and was chosen for additional 3D refinements yielding a 80S ribosome map at an average resolution of 2.39 Å. Particle Subtraction masking either the 40S or the 60S ribosomal subunit granted a separate map of each subunit. The final 40S and 60S maps reached an average resolution of 2.96 Å and 2.44 Å, respectively.

### CONFORMATIONAL HETEROGENEITY ANALYSIS

CryoDRGN (v 1.1.0) (*21*) was used to identify and analyze the ensemble of conformational states present in the non-translating ribosome dataset. Particle poses and CTF data were parsed from the cryoSPARC refinement job output files "particle.cs" using the cryoDRGN utilities parse_pose_csparc and parse_ctf_csparc. Particles were then downsampled from an original box size of 512 pixels to 256 pixels and 128 pixels. Next, variational auto-encoder (VAE) neural network training was performed using the 128-pixel particle set. The default network architecture was used with the number of latent dimensions set to 8 and the VAE was trained for 50 epochs. Outlier particles were identified and removed using GMM (Gaussian Mixture Model) clustering using cryoDRGN utilities implemented in a Jupyter notebook, and a second round of VAE training was performed with the 256-pixel dataset and a larger (1024x3) network model. CryoDRGN reconstructions associated with the 1st and 2nd principal components (PC1 and PC2) were analyzed in ChimeraX (v. 1.4). CryoDRGN training and analysis was performed similarly on the set of ribosomal particles used for the high-resolution reconstruction (consensus map) and for the subset of ~73,000 particles that possessed H69-occluded tRNA binding site, as visualized using cryoSPARC 3D Variability Analysis (focused refinement). See supplementary figure 7 and movies 4, 5, and 6.

### MODEL BUILDING

Model building was done using the software COOT (*49*). Due to the unavailability of a *Schizosaccharomyces pombe* (*S. pombe*) 80S ribosome structure, the initial model was generated by fitting a previously published *Saccharomyces cerevisiae* 80S ribosome model (PDB: 6Z6J) into the cryo-EM reconstruction using UCSF ChimeraX (*50*). Protein and RNA densities were adjusted based on either sequence alignment of all ribosomal proteins and RNA chains between *S. cerevisiae* and *S. pombe* or using AlphaFold models of the *S. pombe* ribosomal proteins. Manual inspection and adjustments of the model were performed in COOT. For the non-translating ribosome, the conformational change in H69 was manually adjusted in COOT. The final model underwent real-space refinement in PHENIX (*51*) against the sharpened electron density map, with the side chain rotamer and Ramachandran restraints options switched on. FSC calculation was reported using the gold standard approach (0.143 threshold) and was validated using the function "map versus model" in PHENIX. All cryo-EM reconstructions and model renderings were generated using ChimeraX (*50*).

### PLATE VIABILITY ASSAY

Wild-type (WT), Δ*dnm1*, and Δ*cpc2* (RACK1) *S. pombe* cells were cultured in YES medium or EMM supplemented with either 2% or 0.5% glucose (w/v) until they reached $OD_{600}$ = 1.0. Subsequently, series of 5-fold cell culture dilutions were prepared. 3 μL droplets of each dilution were deposited onto solid media plates and incubated at 30°C for a duration of 7 days. Images of the plates were taken on days 3 and 7 to monitor growth progression.

### SAMPLE PREPARATION FOR CRYO-ET

Cu 200mesh R2/2 (Quantifoil) grids rendered hydrophilic by plasma cleaning for 30s in a 75%/25% Ar/$O_2$ mixture with a 1070 plasma cleaner (Fischione) were used for plunge-freezing *S. pombe* cells using an EM GP2 plunger (Leica) with the environmental chamber set to 23°C and 100% humidity. The cells were either grown for 1 day in YES medium or for up to 7 days in EMM supplemented with 0.5% glucose (w/v), then diluted right before plunge-freezing to $OD_{600}$ = 0.6 with YES or EMM containing no glucose, respectively. Four µL of the diluted cell suspension were applied to the carbon foil side of the grids which were then blotted for 1 s from the copper mesh side with 597 blotting paper (Whatman) using the blotting paper contact sensing function and 1.3 mm additional move, then immediately plunge-frozen in liquid ethane.

### CRYO-FIB MILLING FOR LAMELLA PREPARATION

Vitrified grids were clipped in CryoFIB Autogrids (ThermoScientific, ref. 1205101) in cold nitrogen vapor, then loaded in an Aquilos DualBeam cryoFIB/SEM (ThermoScientific) operated using the software xT (ThermoScientific). The temperature of both the stage and the anti-contaminating shield were held at -180 °C. The grids were inspected using the SEM operated at 5 kV and 13 pA, then coated with organic Pt (trimethyl (methylcyclopentadienyl) platinum (IV)) for 13 s using the in-built gas injection system (GIS) and finally sputter-coated with metallic Pt for 20 s using a current intensity of 30 mA. Cell monolayers located within 600 µm of the center of the grid were targeted for cryo-FIB milling. Milling was performed using the ion beam operated at 30 kV and currents decreasing from 300 to 30 pA. At 30 pA, lamella thickness was thinned to < 200 nm. During the final stage of lamella milling, the stage was rotated by +0.5° and the top of each lamella was further polished to smoothen thickness variations and limit curtaining artifacts.

### CRYO-ET DATA COLLECTION IN GLUCOSE-DEPLETED WT CELLS

Grid screening and data acquisition were performed on a Titan Krios TEM G3i (ThermoScientific) operated at 300 kV and equipped with a Bioquantum energy filter and a K3 direct electron detector (Gatan) operated in zero-loss mode. On each grid, lamellae were mapped with a pixel size of 28 Å, -100 µm defocus, a 70 µm objective aperture and a 30 eV energy slit. 125 mitochondria were mapped for tilt series (TS) acquisition in SerialEM (*52*) following an established protocol (*53*). Data was acquired at a nominal magnification of 42,000x resulting in a calibrated pixel size of 2.075 Å on the camera. A 50 µm C2 and a 70 µm objective aperture were inserted and the width of the energy filter slit was set to 10 eV. The TEM was operated in nanoprobe mode at spot size 7. TS were acquired using dose-symmetric tilt-scheme (*53*) with 2° tilt increment with tilt angles ranging from -63° to +45° centered on lamella pretilt (≈-9°). Movies were acquired in counting mode over 270 ms exposure time with a total accumulated dose of ~2.23 $e^-/Å^2$ per movie (~15 $e^-$/px/s over an empty area on the camera level) and saved in the TIF file format. 9 frames were saved in each raw tilt image. The accumulated dose of each TS amounted to 120 $e^-/Å^2$. The target defocus was set from -2 µm to -4 µm with 0.5 µm steps between TS. A total of 125 TS was acquired in cells harvested at day 7 of growth in EMM + 0.5% glucose in two overnight imaging sessions using the same microscope and parameters. 53 TS were short-listed for sub-tomogram averaging (STA) based on high dose rate, low lamella thickness and absence of contamination.

### CRYO-ET DATA PROCESSING AND SUB-TOMOGRAM AVERAGING

The cryo-ET data processing and sub-tomogram averaging workflows are shown in fig. S11. Cryo-EM movies were motion-corrected with MotionCor2 (*54*). The alignment of the 53 TS selected for high-resolution STA on membrane-associated ribosomes was performed via semi-automatic tracking of Pt fiducials populating lamellae surfaces using eTOMO (*55*). 3D-CTF correction and tomogram reconstruction were performed using novaCTF (*56*). To initialize STA in all 53 TS, starting positions and orientations of sub-tomograms were generated by manually fitting mitochondria with spheres using IMOD (*57*), and converting these spheres into series of vectors orientated normally to the spheres and separated from each other by 3 px at binning 8 (≈49.8 Å) resulting in 2,091,052 starting positions. Ten rounds of alignment using a cylindrical cross-correlation mask (CC-mask) elongated along the local Z axes (height: 660 Å, radius: 50 Å) generated with Dynamo (*58*) (https://doi.org/10.1016/j.jsb.2011.12.017) locked sub-tomogram positions on the outer mitochondrial membrane (OMM). Particles that diverged away from the OMM were filtered out using the geometry-based procedure described in (*53*) or removed based on visual feedback using the Chimera plugin "Place Object" (*59*). Subsequent rounds of alignment using a wider cylindrical CC-mask (height and radius 166 Å) and a smooth spherical alignment mask (radius: 190 Å; fall-off: 3 px) caused the convergence of sub-tomogram positions on OMM-associated ribosomes. Filtering out positions located closer than 100 Å from one another resulted in a motive list of 42,502 positions in the dataset of 53 TS (53TS). Two TS were short-listed to generate a reference density map for STA in incrementally larger datasets (i.e. 6TS, then 53TS). In the smallest dataset (2TS), consecutive rounds of particle alignment with spherical masks, low-pass filtering cutoff values set always lower than the spatial frequency of a Fourier shell correlation (FSC) score of 0.5 to prevent overfitting, and particle cleaning based on the cross-correlation scores of each particle with the reference density map (CC-cleaning) were performed until obtaining a map with a resolution of ≈25 Å (FSC score of 0.143) based on 541 particles. This reference was used to align ribosomes in the 6TS dataset with similar STA parameters. Consecutive rounds of alignment and CC-cleaning in the 6TS dataset provided a map with a resolution of ≈20 Å based on 701 particles. Using this reference for rounds of alignment and CC-cleaning in the 53TS dataset provided a map with a resolution of ≈15Å based on 10,192 particles. To reach higher resolution, the coordinates of these particles were imported in Warp (*60*) following conversion of the motive list in .star format using in-house scripts. The TS and alignment files were also imported in Warp using in-house scripts (https://git.embl.de/gemin/ribosonmitos). 53 tomograms were reconstructed from which the 10,192 sub-tomograms were re-extracted and loaded in the software RELION (*61*) for per-particle CTF estimation and 3D alignment. The tomograms were then ported to the software M (*62*) for consecutive rounds of alignment, denoising and image & volume warping resulting in a density map with a resolution of 11.4 Å (selected options: Image warp grid 5x5, Particle poses, Stage angles and Volume warp grid 8x8x2x10).

### INTERFACES OF NEIGHBORING RIBOSOMES WITHIN PENTAMERS

The relative position and orientation of wild-type neighboring ribosomes within pentamers (shown in Fig. 3) were derived from the position and orientation of individual ribosomes in pentamers. Pentamer template recognition was first performed on the full motive list (10,192 particles) to extract the coordinates of ribosomes engaging in pentamer formation (github/Nmer_parameter_find.m & SM_find_pentamer_normal.m). Ribosomes in each extracted pentamer were then sorted clock-wise and re-numbered from 1 to 5. Translations relating neighboring ribosomes were derived from vectors $\mathbf{v_n}$-$\mathbf{v_m}$ with $\mathbf{v_n}$ the cartesian coordinates of ribosome n and {n,m} taking values: {1,2},{2,3},{3,4},{4,5},{5,1}. Differences in orientation between neighboring ribosomes were estimated by rotation matrices $\boldsymbol{R_{n,m}} = \boldsymbol{R_n}^{-1}\boldsymbol{R_m}$ with $\mathbf{R_n}$ the rotation matrix transforming $\mathbf{u_z}$(0,0,1) into $\mathbf{u_{z,n}}$ the local "upward" unit vector of ribosome n. Each $\mathbf{R_{n,m}}$ was converted to a quaternion (github/euler2quat.m) $\mathbf{q_{n,m}}$. To visualize the interaction mode of ribosomes contacting each other within pentamers, we selected the 10% closest pairs of ribosomes (n=62 out of 625 pairs) and computed the average translation and rotation describing this ensemble, $<\mathbf{v_n}-\mathbf{v_m}>_{10\%}$ and $<\mathbf{q_{n,m}}>_{10\%}$ using Markley's method (github/avg_quaternion_markley.m). To display this interaction in Fig. 3, we transformed the cartesian coordinates $\mathbf{P_1}$(x,y,z) of each atom of our PDB structure of hibernating ribosome into $\mathbf{P_2} = <\mathbf{q_{n,m}}>_{10\%}\mathbf{P} + <\mathbf{v_n}-\mathbf{v_m}>_{10\%}$ and displayed both structures using ChimeraX (*50*).

### CRYO-ET DATA COLLECTION AND PROCESSING IN MUTANT YEAST STRAINS

Cryo-ET data of Δ*dnm1* and Δ*cpc2* cells harvested at day 7 of growth in EMM + 0.5% glucose (w/v) were acquired using a Titan Krios TEM G4 (ThermoScientific) operated at 300 kV and equipped with a Selectris X energy filter and a Falcon 4i direct electron detector (ThermoScientific) operated in zero-loss mode. Lamellae screening was performed in similar conditions as for fully starved cells. Tilt series acquisitions were performed using the parameters reported in table S3. As for processing, TS were automatically aligned and reconstructed as tomograms using AreTomo (*63*); ribosome picking was performed by gaussian fitting using in-house python scripts (github/BlobPicking.py); ribosome picks were cleaned manually using IMOD; particle orientations were estimated using the sub-stack averaging pipeline SUSAN (*64*). Quick particle alignment in SUSAN resulted in preliminary structures of cytosolic ribosomes resolved at 45 Å resolution in Δ*cpc2* cells and 33 Å resolution in Δ*dnm1* cells. This enabled aligning and orienting ribosome models in 3D renderings of the cytosol of these cells displayed in Fig. 4 and 5.

### 3D RENDERING OF MITOCHONDRIA

Models of mitochondrial membranes displayed in figures 2, 4 and 5 were segmented in IMOD using the plugin "Drawing Tools", then meshed and imported in Blender (https://www.blender.org/) as .obj files. Ribosomes were placed in the 3D rendering layout based on motive lists converted into .star files (github/em2bdStar.m and .blend files) using the plugin Molecular Nodes (*65*).

### SUPPLEMENTARY FIGURES:

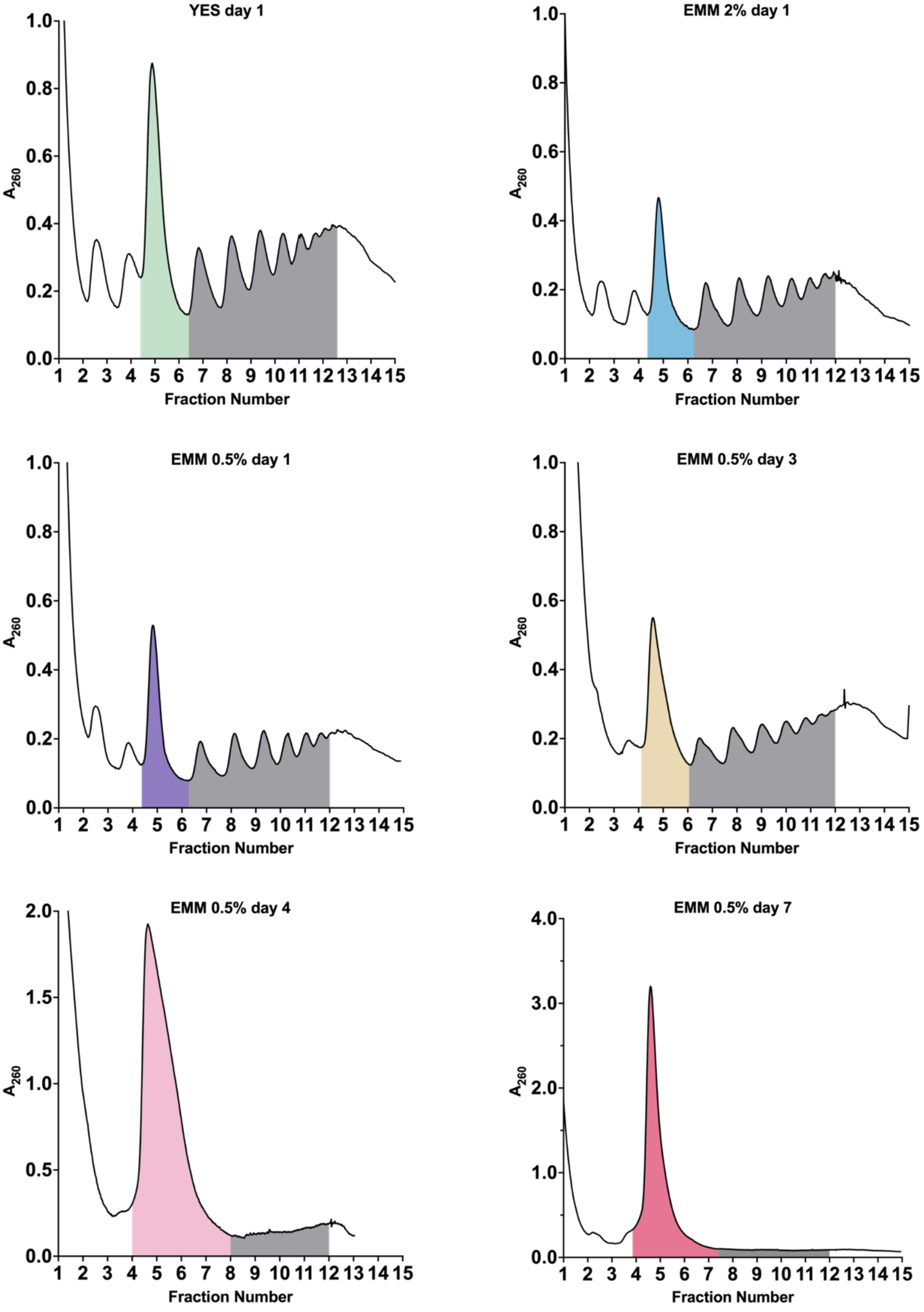

**Fig. S1. Polysome profiles for *S. pombe* grown under low and high glucose conditions.**

Polysome profiles obtained from sucrose density gradients of cell lysates from *S. pombe* cultures grown in either YES or EMM media supplemented with 2% or 0.5% glucose (w/v). Cells were harvested as time-lapse at days 1, 3, 4 or 7 for analysis. Areas under the 80S peaks are highlighted using the same colors as in Fig. 1C, and areas under polysome fractions are highlighted in gray.

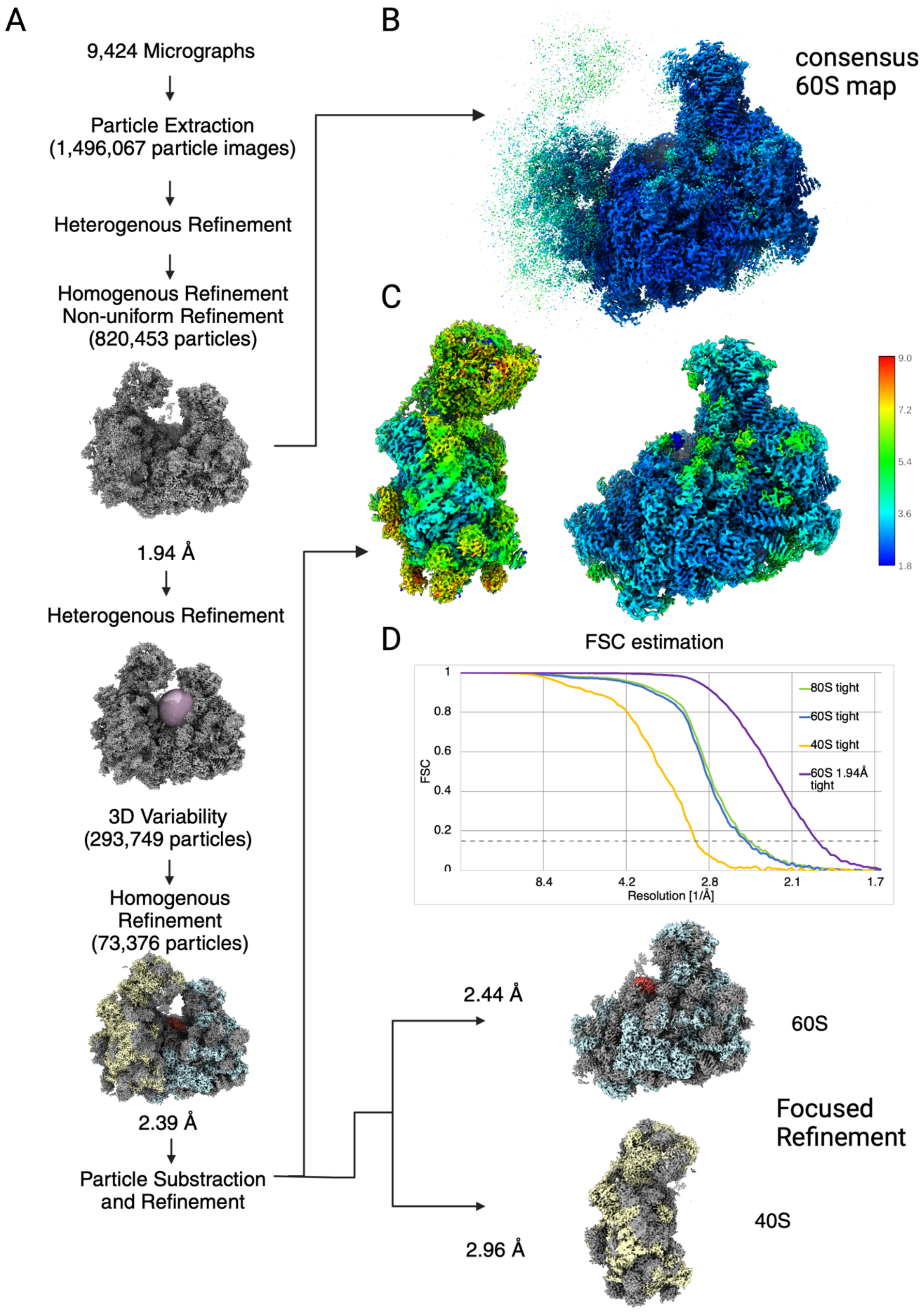
A
9,424 Micrographs
Particle Extraction
(1,496,067 particle images)
Heterogenous Refinement
Homogenous Refinement
Non-uniform Refinement
(820,453 particles)
1.94 Å
Heterogenous Refinement
3D Variability
(293,749 particles)
Homogenous
Refinement
(73,376 particles)
2.39 Å
Particle Substraction
and Refinement
B
consensus
60S map
C
9.0
7.2
5.4
3.6
1.8
D
FSC estimation
FSC
Resolution [1/Å]
1
0.8
0.6
0.4
0.2
0
8.4
4.2
2.8
2.1
1.7
80S tight
60S tight
40S tight
60S 1.94Å
tight
2.44 Å
60S
Focused
Refinement
2.96 Å
40S

**Fig. S2. Processing scheme of the single-particle analysis of non-translating ribosomes.**

(A) Particles were picked from motion-corrected and dose-weighted micrographs in cryoSPARC. Heterogenous refinement followed by Homogenous and Non-uniform refinements resulted in a consensus map of non-translating 80S ribosomes. 3D variability analysis using a binary mask around the PTC was performed to select particles exhibiting the best-resolved conformational change in H69 (colored red). Homogenous refinement of selected particles was followed by particle subtraction to obtain high-resolution structures of both the 40S and 60S ribosome subunits.

(B) and (C) 3D representations of local resolution calculated for the final 80S, 60S and 40S ribosome maps.

(D) Fourier shell correlation plot for the final maps.

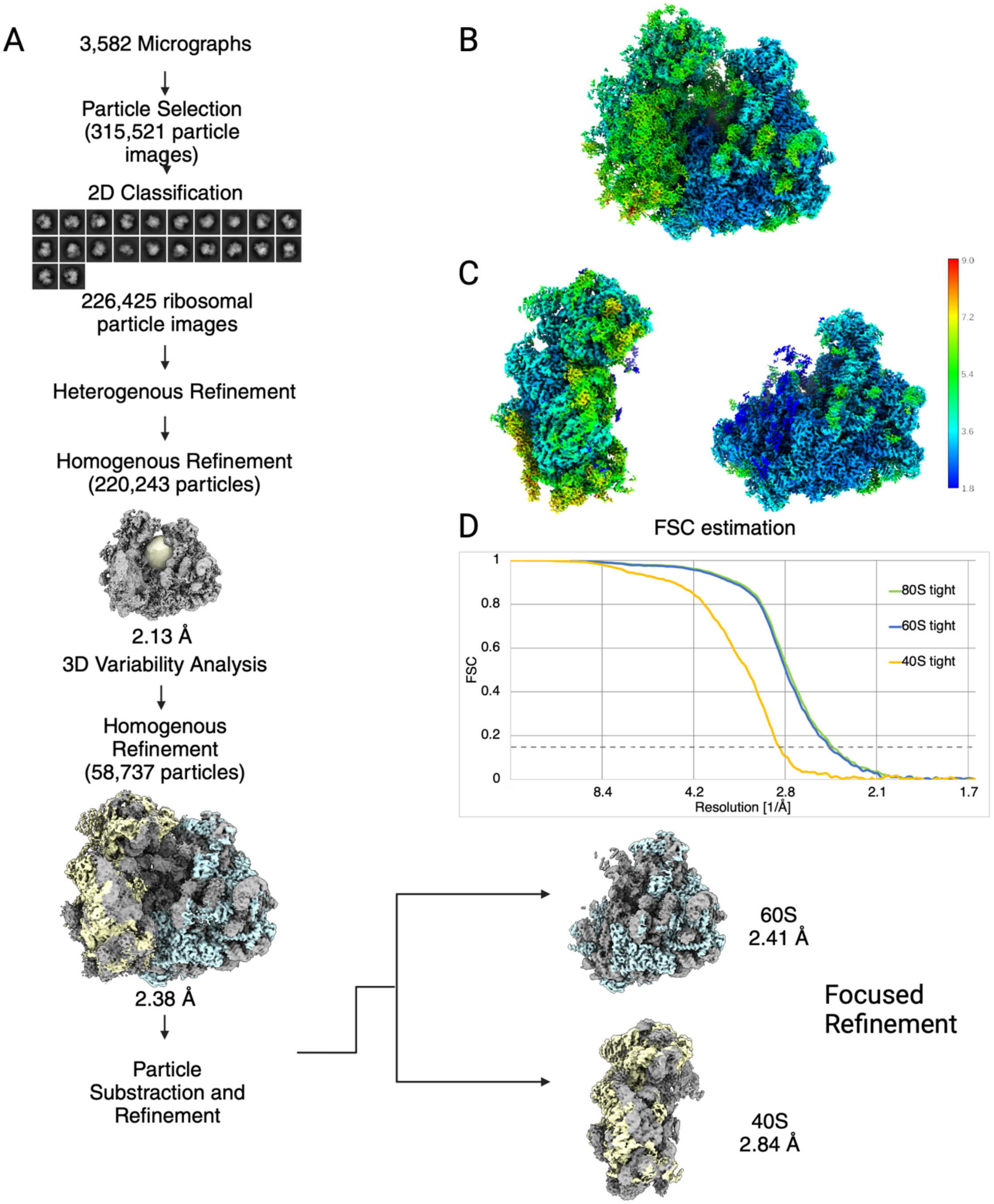


**Fig. S3. Processing scheme of the single-particle analysis of translating ribosomes.**

(A) Particles were picked from motion-corrected and dose-weighted micrographs using blob picker in cryoSPARC. 2D-classification followed by Heterogenous and Homogenous

Refinement resulted in the initial 80S ribosome. 3D variability analysis was performed to obtain an 80S ribosome using the particles exhibiting the best-resolved tRNA densities. To improve resolution of 40S and 60S subunits, particle extraction was performed using binary masks for 60S and 40S subunits, respectively.

(B) and (C) 3D representations of local resolution calculated for the final 80S, 60S and 40S ribosome maps.

(D) Fourier shell Correlation plot for the final maps.

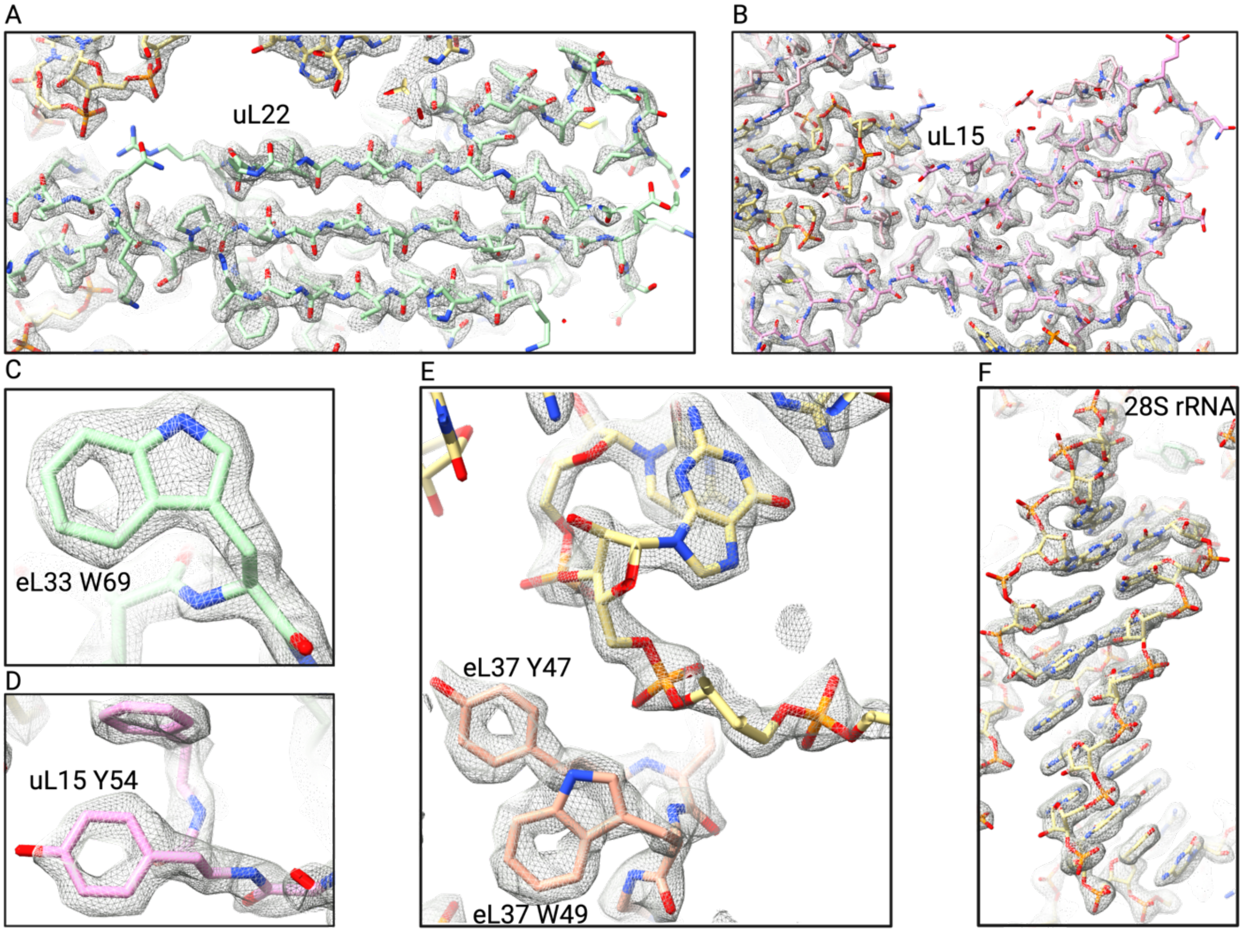


**Fig. S4. Close-up on densities of ribosomal protein and rRNA reconstructions.**

Ribosomal proteins and rRNA shown as sticks with overlaid cryo-EM maps of *S. pombe* non-translating ribosome shown as gray mesh.

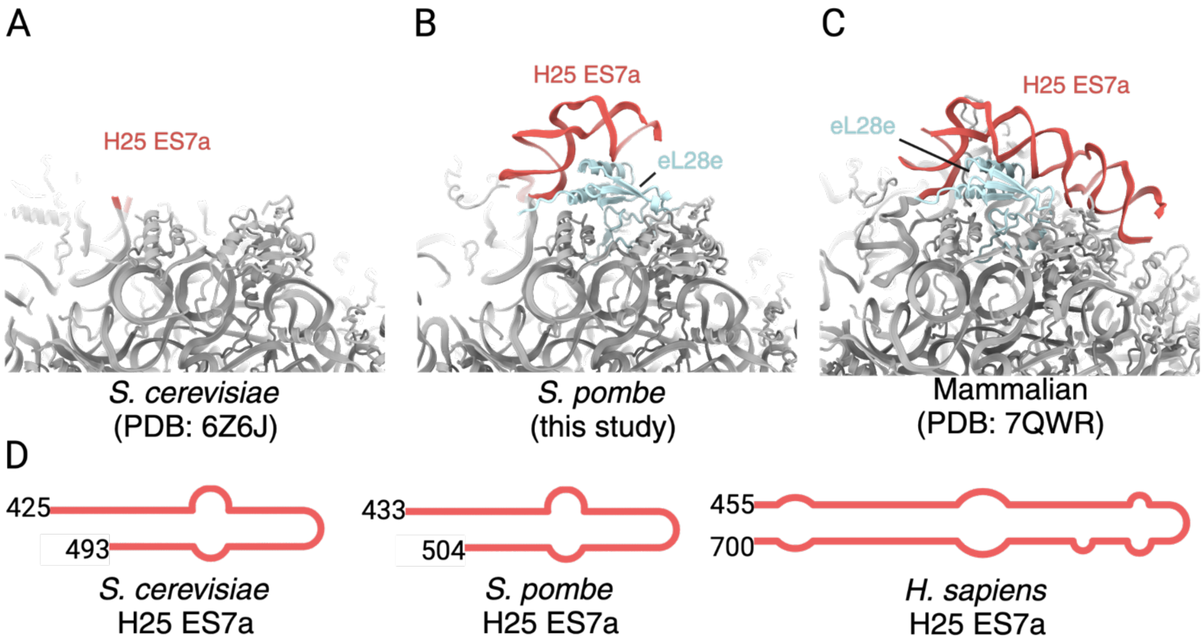


**Fig. S5. L28e and H25 Expansion Segment 7a comparison.**

(A) Cryo-EM ribosome structure of a *Saccharomyces cerevisiae* ribosome (PDB: 6Z6J) lacking ribosomal protein eL28e.

(B) Cryo-EM ribosome structure of a *Schizosaccharomyces pombe* ribosome showing the presence of ribosomal protein eL28e and subsequent ordering of H25 ES7a.

(C) Cryo-EM ribosome structure of a mammalian ribosome (PDB: 7QWR) showing the presence of ribosomal protein eL28e and subsequent ordering of H25 ES7a.

(D) Secondary structure representations of H25 ES7a in *S. cerevisiae*, *S. pombe* and *H. sapiens*.

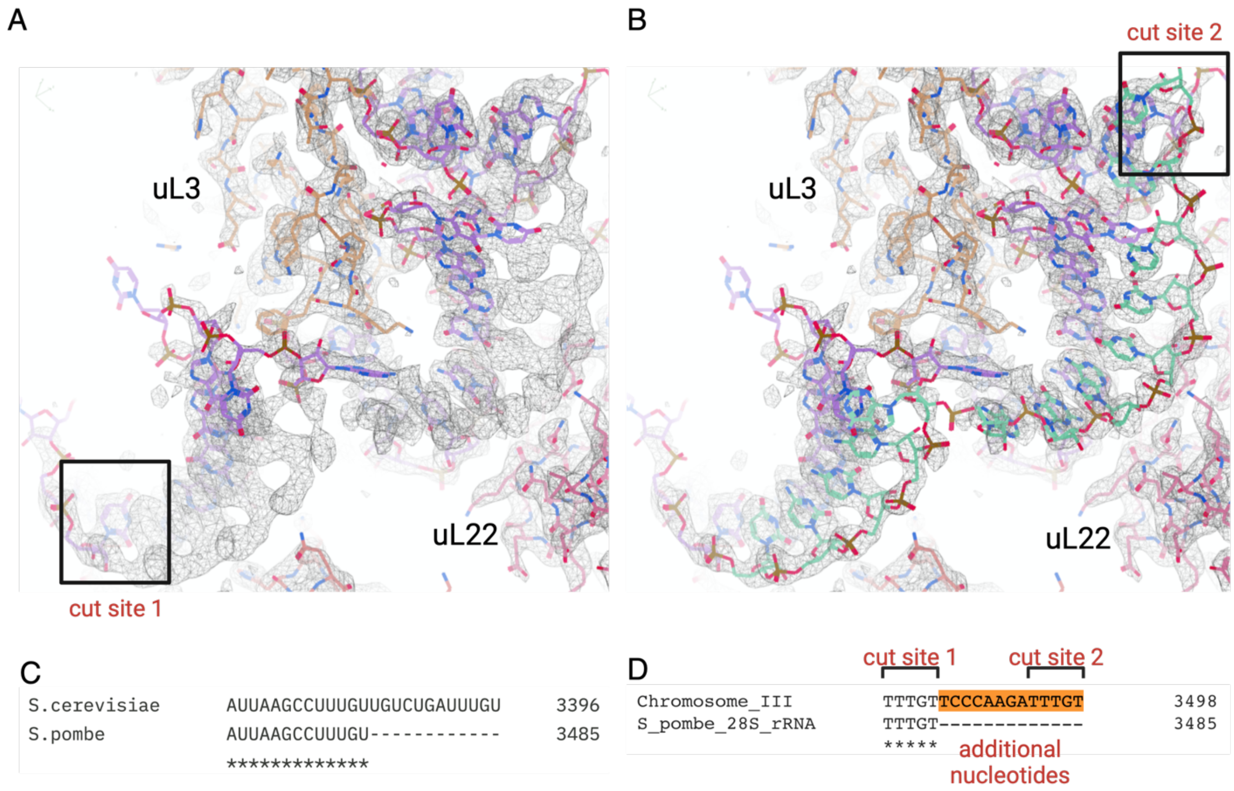


**Fig. S6.** ***S. pombe*** **28S rRNA non-annotated nucleotides resolved in the cryo-EM map.**

(A) Cryo-EM map of the *S. pombe* ribosome showing additional density at 3’ end of 28S rRNA and overlaid with the preexisting model of 28S rRNA. The last nucleotide of previously annotated 28S rRNA sequence is boxed.

(B) Cryo-EM electron density map of *S. pombe* ribosome and its model supplemented with 13 additional nucleotides based on the sequence of chromosome III. The last missing nucleotide of previously annotated 28S rRNA sequence is boxed.

(C) Sequence alignment of *S. cerevisiae* 25S rRNA and *S. pombe* 28S rRNA.

(D) Sequence alignment of *S. pombe* chromosome III and 28S rRNA. Repetitive nucleotide sequence shown in brackets. The 13 nucleotides missing from 28S rRNA sequence are highlighted in orange.

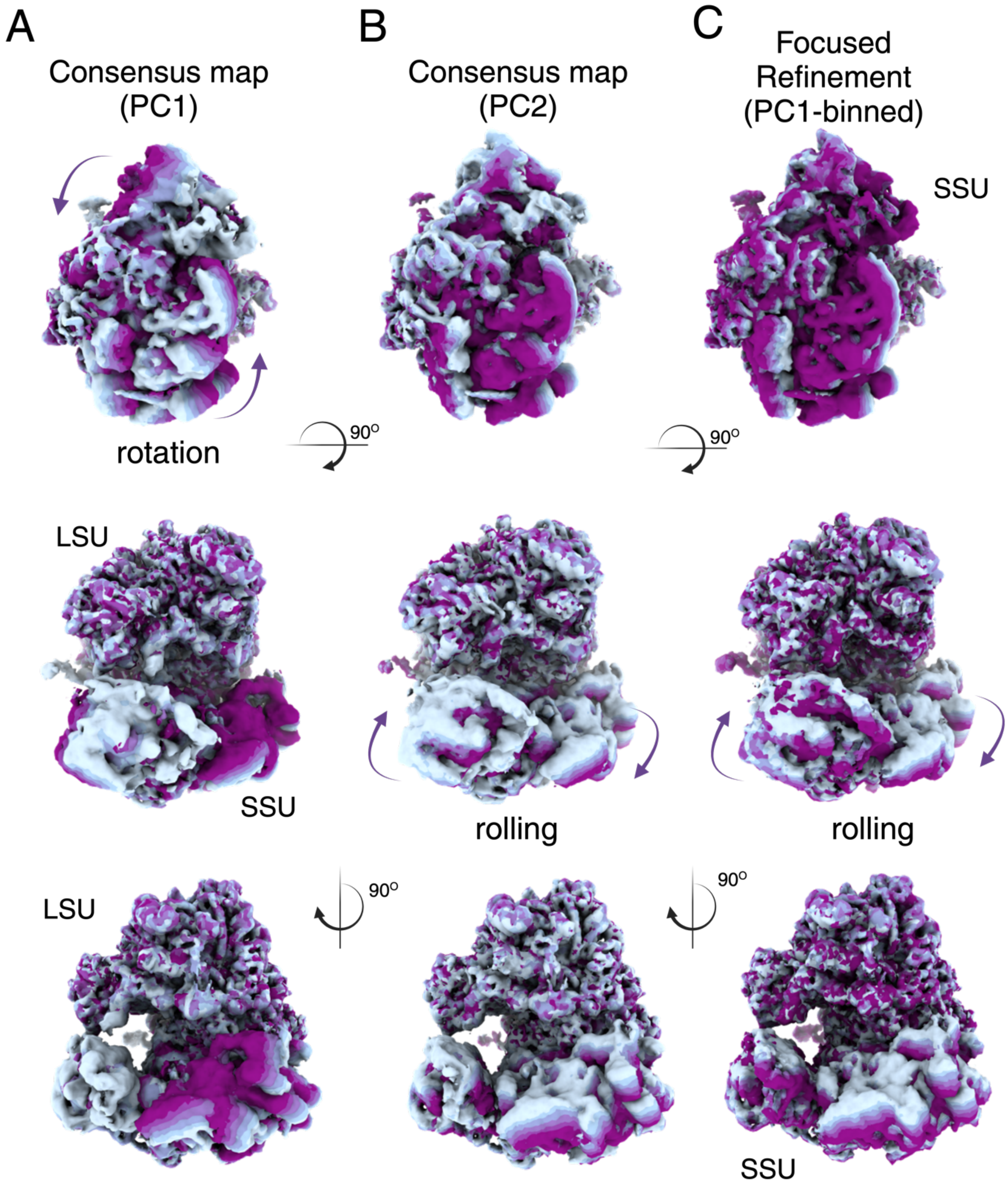


**Fig. S7. Probing conformational dynamics of the SSU in the non-translating ribosome using CryoDRGN.**

(A) and (B) CryoDRGN was applied to the dataset from which the high-resolution consensus map was derived to interrogate SSU conformational space. Following principal component analysis (PCA), we identified two main conformational changes: the rotation or rolling of the SSU relatively to the LSU (principal components PC1 and PC2 shown with magenta arrows, respectively).

(C) CryoDRGN applied to particles in which H69 occludes the PTC (i.e. the dataset used for the focused refinement in fig. S2). This analysis highlights a rolling movement similar to PC2 shown in (B). Also see Movies S4-6.

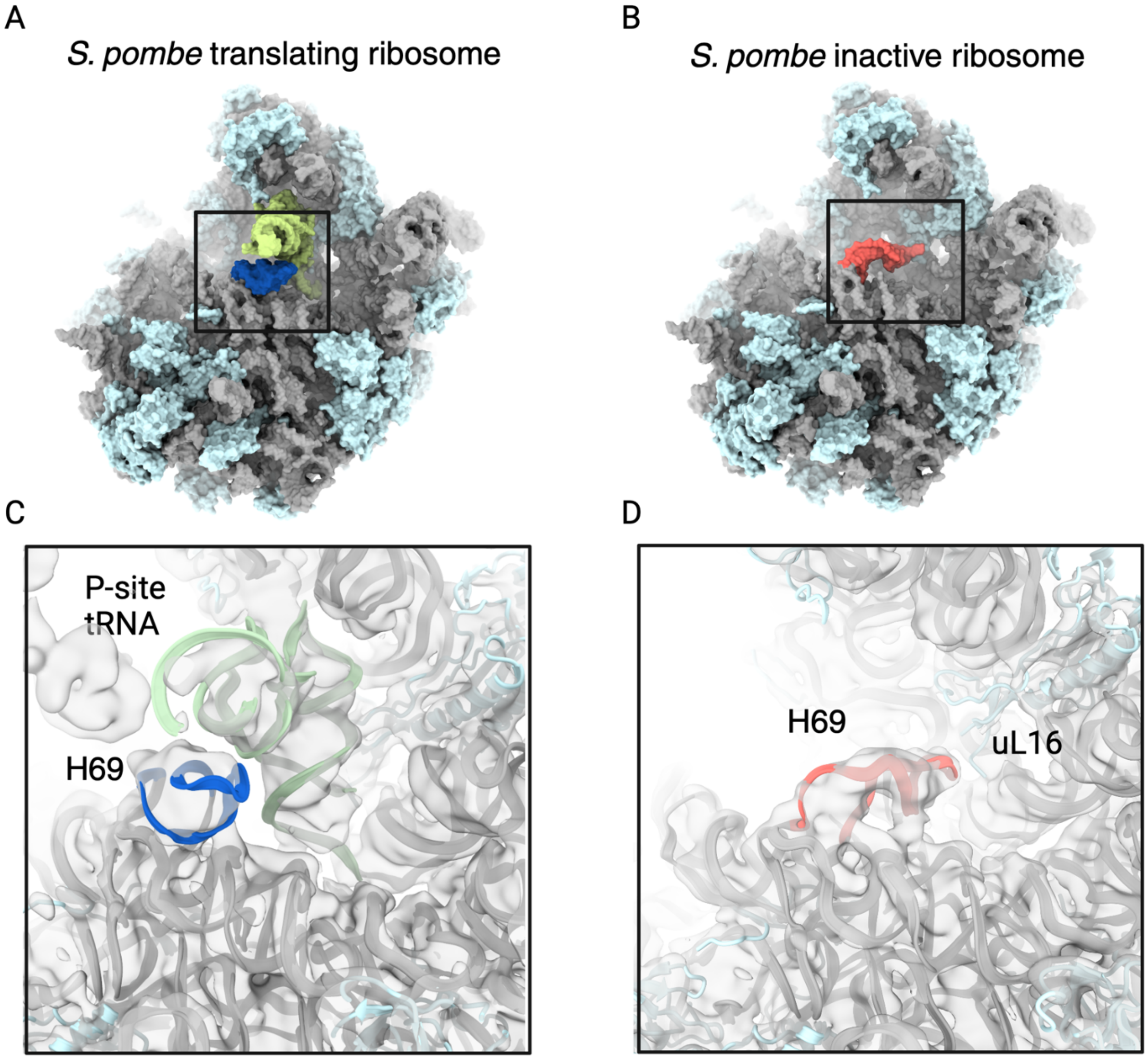


**Fig. S8. Cryo-EM structure of both the translating and the inactive *S. pombe* ribosome.**

(A) Atomic model of the LSU of the translating ribosome shown as surface. Helix H69 is highlighted in royal blue. tRNA, rRNA and r-proteins are colored green, gray and cyan, respectively.

(B) Atomic model of the LSU of the inactive ribosome shown as surface. Helix H69 is highlighted in red.

(C) Close-up on the area boxed in (A) shown as a ribbons and overlaid with the EM density map filtered to 6 Å resolution and shown as a transparent surface.

(D) Close-up on the area boxed in (B).

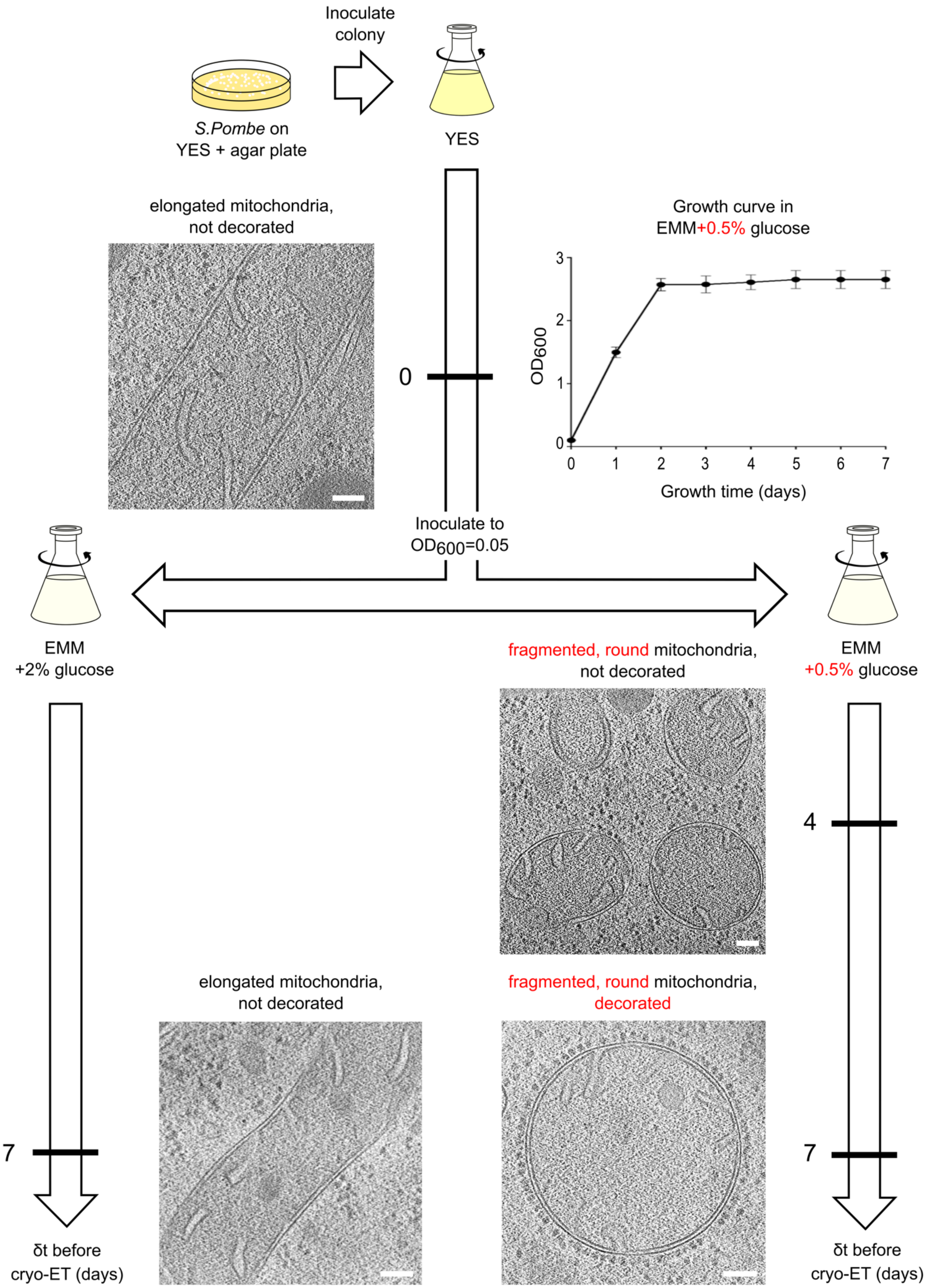
Inoculate
colony
S.Pombe on
YES + agar plate
YES
elongated mitochondria,
not decorated
0
Growth curve in
EMM+0.5% glucose
OD600
Growth time (days)
Inoculate to
OD600=0.05
EMM
+2% glucose
EMM
+0.5% glucose
fragmented, round mitochondria,
not decorated
4
elongated mitochondria,
not decorated
fragmented, round mitochondria,
decorated
7
7
δt before
cryo-ET (days)
δt before
cryo-ET (days)

**Fig. S9. Time-course *in situ* cryo-ET for WT *S. pombe* cells growing under different glucose conditions.**

Following inoculation of *S. pombe* cells grown on a YES+agar plate into a YES liquid culture, then an EMM liquid culture, cells were harvested at day 0 and 7 for EMM + 2% glucose (w/v) and days 0, 4, and 7 for EMM + 0.5% glucose (w/v), then plunge-frozen on cryo-EM grids. $OD_{600}$ measurements were taken at all 7 days of cell growth for n=7 cultures in EMM + 0.5% glucose. Cryo-ET cross sections of representative mitochondria at the indicated time points are shown, highlighting that under low glucose conditions, mitochondria become fragmented after 4 days and decorated with ribosomes at day 7. Scale bars 100 nm.

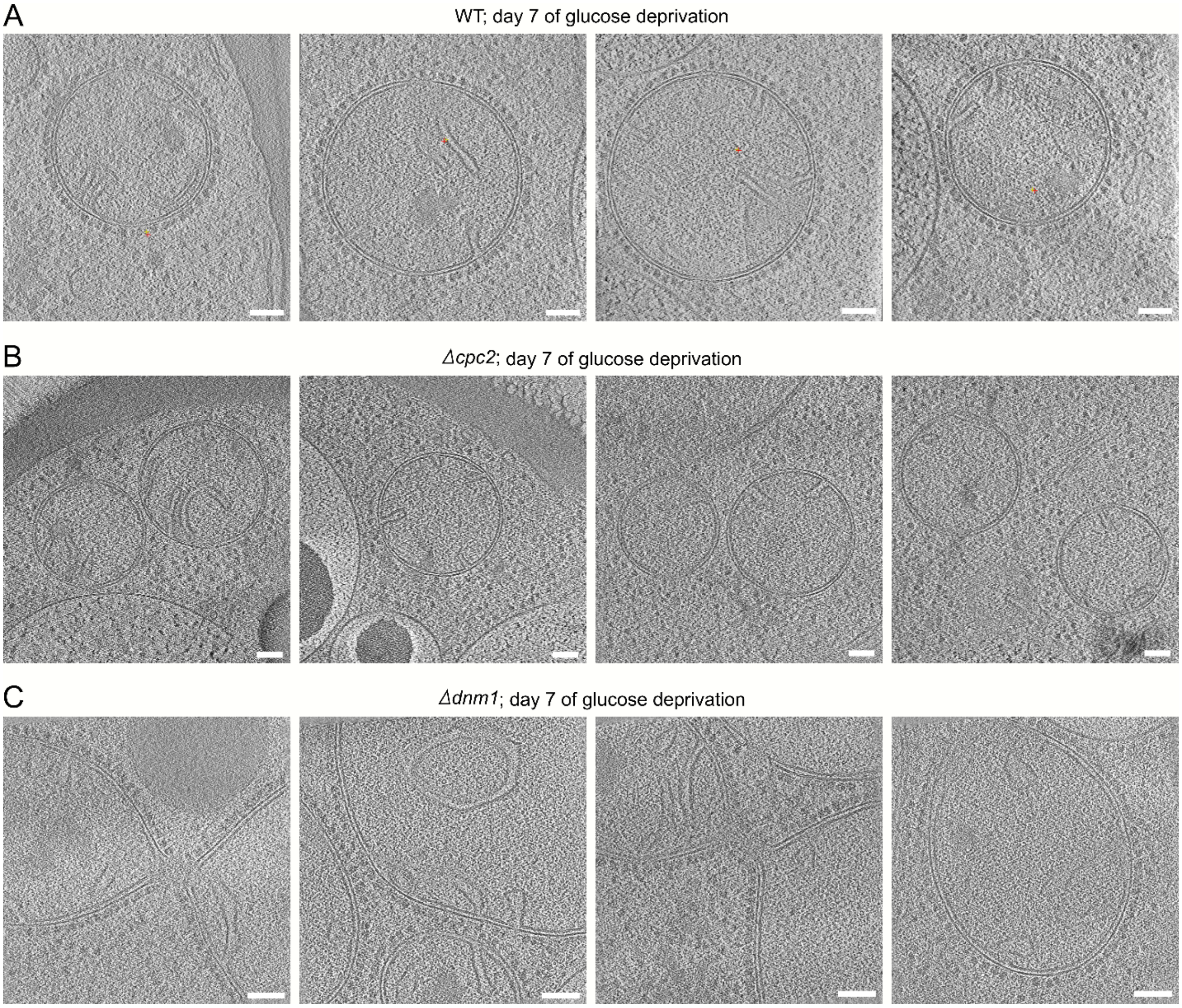


**Fig. S10. In WT or mutant *S. pombe* cells grown for 7 days at low glucose concentration, the vast majority of mitochondria adopt a strain-dependent morphology.**

Representative cryo-ET cross sections of mitochondria at day 7 of glucose depletion in WT *S. pombe* cells (A), in *Δcpc2* cells (B) and in *Δdnm1* cells (C). Scale bars 100 nm.

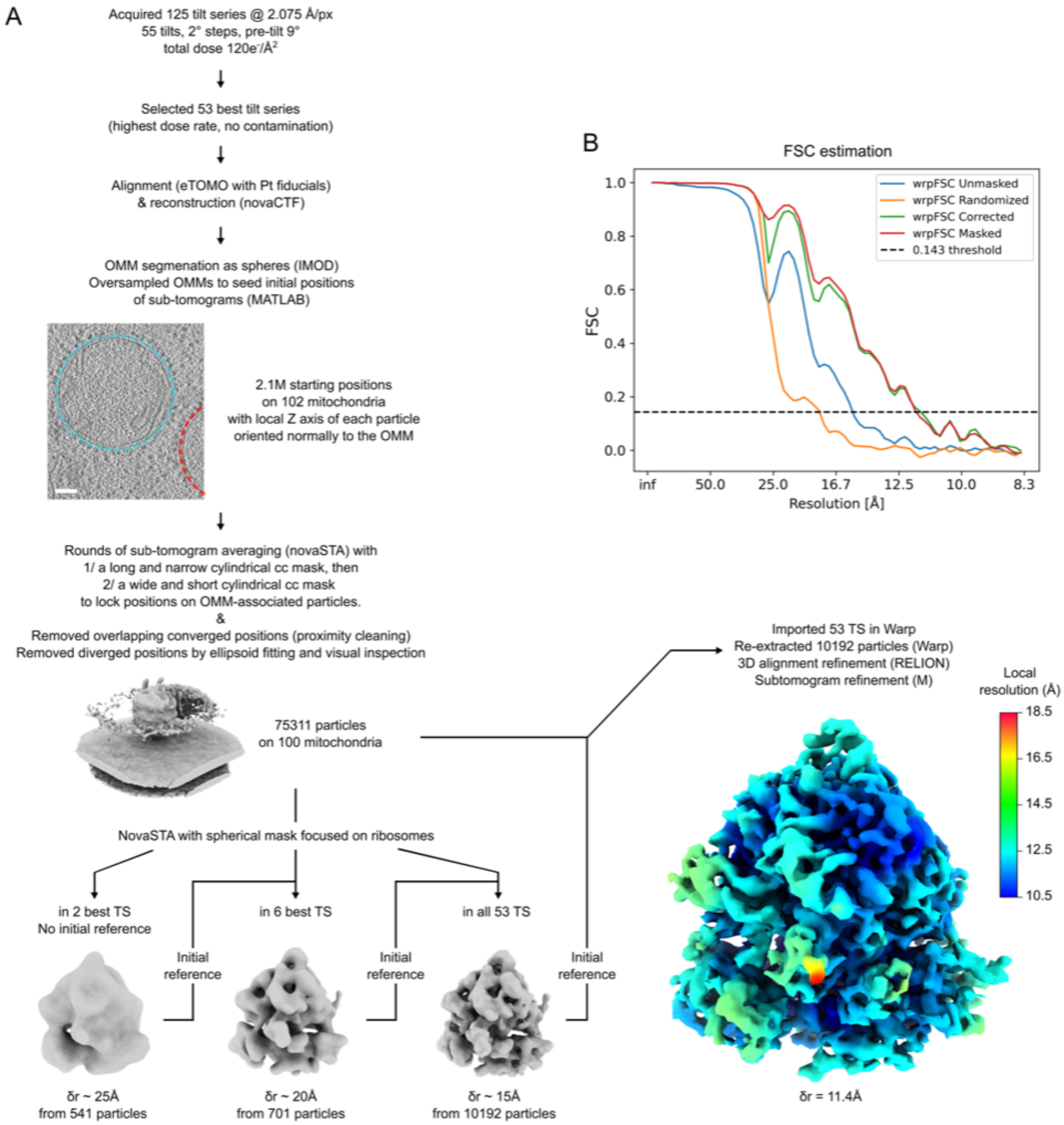


**Fig. S11. Cryo-ET and subtomogram averaging processing scheme**

(A) Flow-chart of the cryo-ET pipeline used on WT *S. pombe* cells: following cell freezing at day 7 of glucose depletion and lamellae milling, dose-symmetric tilt series were acquired using a Titan Krios TEM. The 53 best tilt-series were aligned, reconstructed as tomograms and segmented to initialize subtomogram extraction along the OMM of round-up mitochondria. Consecutive rounds of particle alignment locked sub-tomogram positions on OMM-associated ribosomes. Cleaning steps removed duplicates and particles that were dissimilar to the STA references. Alignment refinement using novaSTA, Warp, RELION and M allowed reconstructing a density map of OMM-associated ribosomes with a resolution of 11.4 Å using 10,192 particles.

(B) Fourier shell correlation (FSC) curve of the cryo-ET reconstruction of the *S. pombe* hibernating ribosome refined using M.

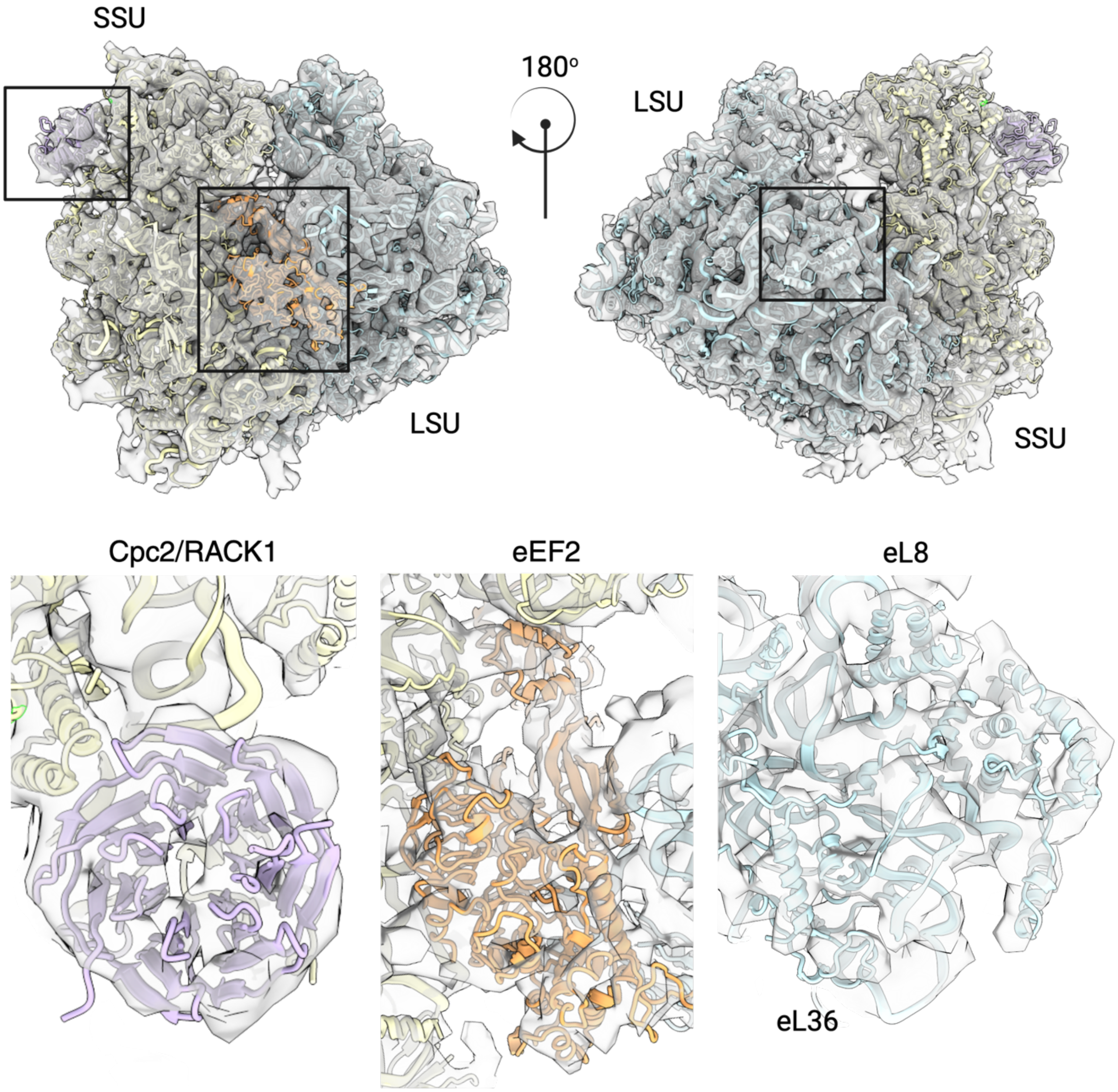


**Fig. S12. *In-situ* cryo-ET structure of the hibernating *S. pombe* ribosome.**

*In-situ* cryo-ET map of the hibernating ribosome with underlying atomic model determined by the high-resolution cryo-EM docked as rigid-body. Close-ups of the boxed areas of Cpc2, eEF2, eL8 are shown in the bottom panels and colored as in the main figures.

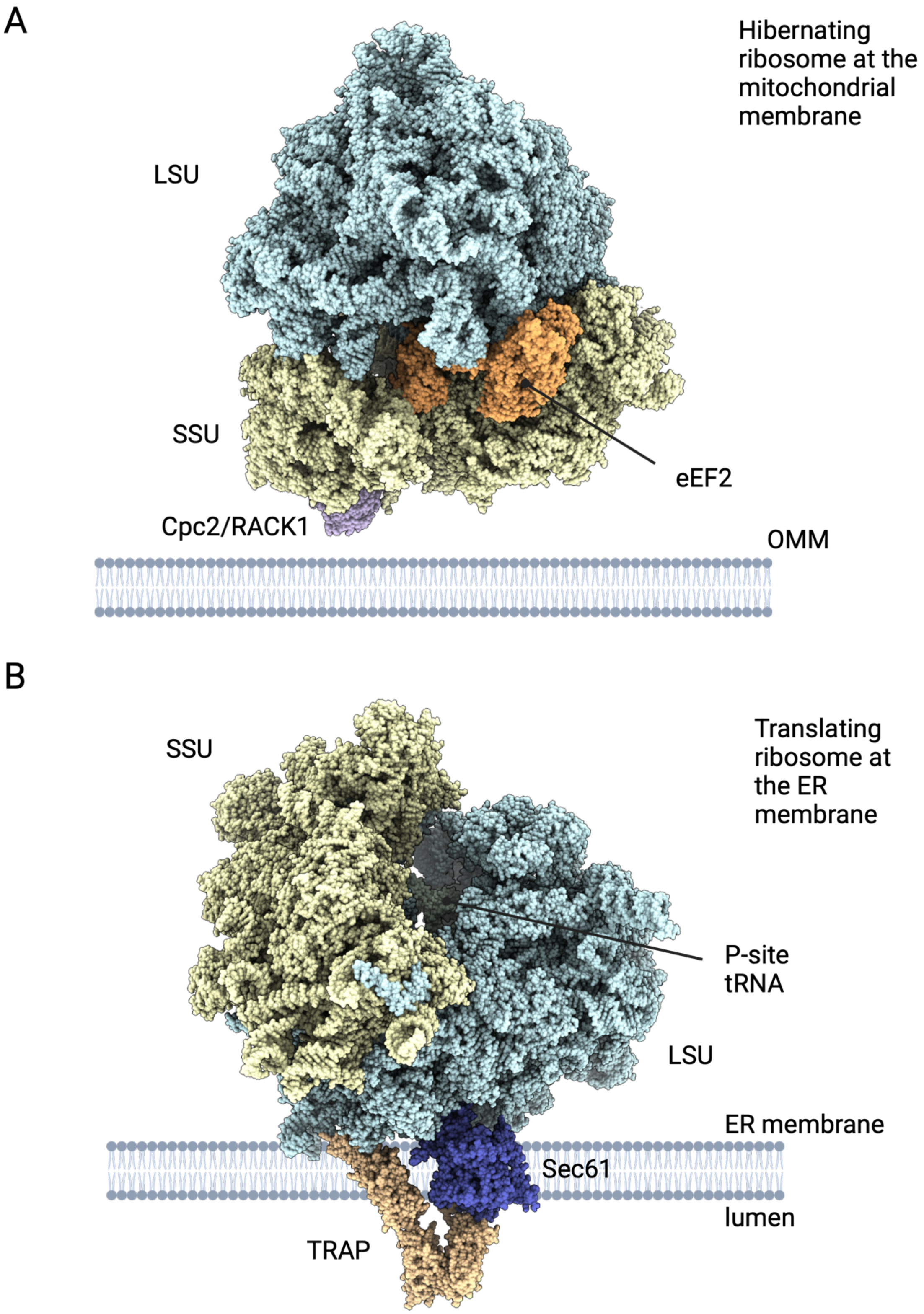
A
Hibernating ribosome at the mitochondrial membrane
LSU
SSU
eEF2
Cpc2/RACK1
OMM
B
SSU
Translating ribosome at the ER membrane
P-site tRNA
LSU
ER membrane
Sec61
lumen
TRAP

**Fig. S13. Comparing an OMM-tethered hibernating ribosome with an ER-docked translating ribosome.**

(A) *In situ* structure of hibernating ribosomes tethered to a mitochondrion. Atoms shown as spheres. LSU and SSU are colored blue and yellow, respectively. eEF2 is colored orange, and Cpc2 is colored purple.

(B) Structure of the ER-docked translating ribosome (PDB: 8BTK). P-site tRNA is colored green, TRAP complex is colored beige, and Sec61 is colored blue.

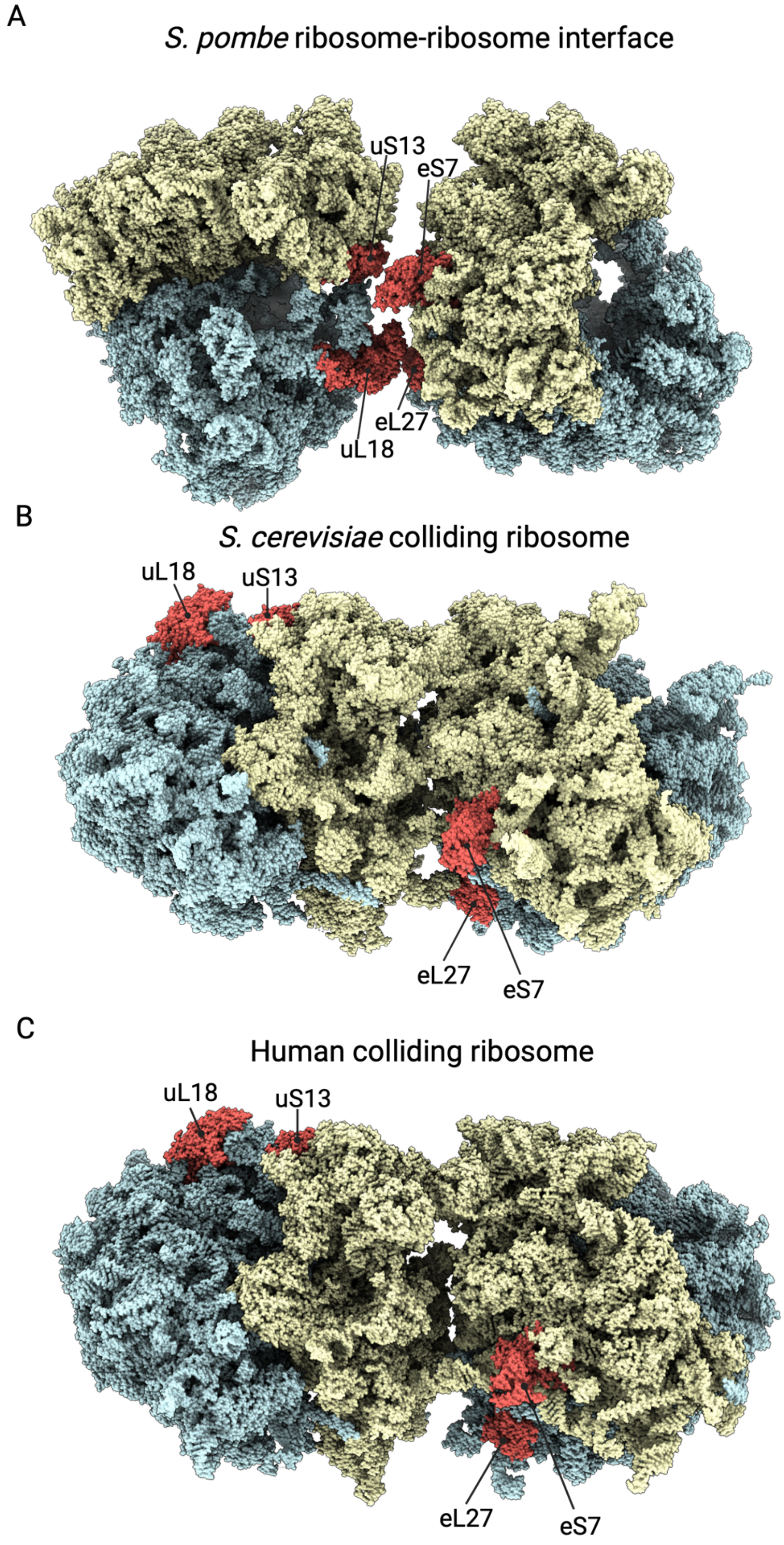
A
S. pombe ribosome-ribosome interface
uS13
eS7
eL27
uL18
B
S. cerevisiae colliding ribosome
uL18
uS13
eL27
eS7
C
Human colliding ribosome
uL18
uS13
eL27
eS7

**Fig. S14. Comparison of *S. pombe* hibernating ribosome-ribosome interface and colliding eukaryotic ribosomes.**

(A) *Schizosaccharomyces pombe* ribosome-ribosome interface at the OMM (this study).

(B) *Saccharomyces cerevisiae* colliding ribosomes (PDB: 6I7O).

**(C)** Human colliding ribosomes (PDB: 7QVP). Large subunit colored cyan, small subunit colored yellow, interacting proteins in *S. pombe* colored in red.

**Table S1. EM data collection and structure refinement statistics.**

| | **Inactive 60S ribosome consensus map** | **Inactive 80S ribosome** | **Translating 80S ribosome** |
|---|---|---|---|
| EMDB code | XXXX | XXXX | XXXX |
| PDB code | YYYY | YYYY | YYYY |
| **Data collection and processing** | | | |
| Nominal magnification | 105,000x | 105,000x | 105,000x |
| Voltage (kV) | 300 | 300 | 300 |
| Electron exposure ($e^{-}$/Å$^2$) | 50 | 50 | 50 |
| Defocus range (µm) | -1.6/-0.6 | -1.6/-0.6 | -1.6/-0.6 |
| Pixel size (Å) | 0.83 | 0.83 | 0.83 |
| Initial particle images (no.) | 1,214,576 | 1,214,576 | 315,521 |
| Final particle images (no.) | 820,453 | 73,376 | 58,737 |
| Map resolution at FSC=0.143 (Å) | 1.94 (60S) | 2.44/2.96 (60S/40S) | 2.27/2.75 (60S/40S) |
| **Structure refinement in PHENIX 1.20.1** | | | |
| Model resolution at FSC=0.5 (Å) | 2.2 | 2.9 | 2.8 |
| $CC_{mask}$ | 0.87 | 0.77 | 0.78 |
| Map sharpening B factor (Å$^2$) | 47.4 (60S) | 43.7/47.6 (60S/40S) | 42.4/46.4 (60S/40S) |
| Model composition | | | |
| Non-hydrogen atoms | 123,483 | 194,864 | 197,314 |
| Protein residues | 6,134 | 10,542 | 10,855 |
| RNA residues | 3,488 | 5,198 | 5,198 |
| B factors min/max/mean (Å$^2$) | | | |
| Protein | 0.00/73.29/30.39 | 0 00/76.07/37 47 | 0.00/65.78/31.57 |
| RNA | 0.00/109.27/40.41 | 0.00/122.76/54.78 | 0.00/105.21/44.85 |
| Ligand | 19.03/88.04/37.31 | 16.71/89.66/43.54 | 5.81/84.22/37.97 |
| RMSD | | | |
| Bond lengths (Å) | 0.003 | 0.002 | 0.002 |
| Bond angles (°) | 0.673 | 0.476 | 0.5 |
| **Validation** | | | |
| MolProbity score | 2.03 | 2.10 | 2.33 |
| Clashscore | 7.38 | 8.79 | 9.15 |
| Poor rotamers (%) | 2.87 | 2.61 | 3.73 |
| Ramachandran plot | | | |
| Favored (%) | 96.03 | 95.38 | 93.80 |
| Allowed (%) | 3.89 | 4.30 | 5.91 |
| Disallowed (%) | 0.08 | 0.12 | 0.29 |

**Table S2. LSU and SSU proteins of the S. pombe ribosome**

| Chain ID | Chain name | PDB entry |
|---|---|---|
| AD | Small ribosomal subunit protein uS2A | Q9Y7L8 |
| AE | Small ribosomal subunit protein eS1 | O94438 |
| AF | Small ribosomal subunit protein uS5 | O74892 |
| AG | Small ribosomal subunit protein uS3 | O60128 |
| AH | Small ribosomal subunit protein uS4 | Q9P4W9 |
| AI | Small ribosomal subunit protein uS5 | Q9P3T6 |
| AJ | Small ribosomal subunit protein eS6B | Q9C0Z7 |
| AK | Small ribosomal subunit protein uS7 | Q10101 |
| AL | Small ribosomal subunit protein eS8 | Q9P7B2 |
| AM | Small ribosomal subunit protein uS4B | P0CT65 |
| AN | Small ribosomal subunit protein eS10B | O13614 |
| AO | Small ribosomal subunit protein uS17B | P0CX48 |
| AP | Small ribosomal subunit protein eS12A | P0CT75 |
| AQ | Small ribosomal subunit protein uS15 | P28189 |
| AR | Small ribosomal subunit protein uS11B | P0CT57 |
| AS | Small ribosomal subunit protein uS19B | Q9UTQ6 |
| AT | Small ribosomal subunit protein uS9B | P0CX52 |
| AU | Small ribosomal subunit protein eS17A | O42984 |
| AV | Small ribosomal subunit protein uS13B | P0CT67 |
| AW | Small ribosomal subunit protein eS19A | P58234 |
| Aa | Small ribosomal subunit protein uS10 | O74893 |
| Ab | Small ribosomal subunit protein eS21 | P05764 |
| Ac | Small ribosomal subunit protein uS8A | P0CT58 |
| Ad | Small ribosomal subunit protein uS12A | P0CT75 |
| Ae | Small ribosomal subunit protein eS24A | O13784 |
| Af | Small ribosomal subunit protein eS25A | P79009 |
| Ag | Small ribosomal subunit protein eS26A | Q9UT56 |
| Ah | Small ribosomal subunit protein | O74330 |
| Ai | Small ribosomal subunit protein eS28A | P0CT79 |
| Aj | Small ribosomal subunit protein uS14 | O74329 |
| Ak | Small ribosomal subunit protein eS30B | P0CT63 |
| Am | Small ribosomal subunit protein RACK1 | Q10281 |
| B0 | Large ribosomal subunit protein eL42 | Q9UTI8 |
| B1 | Large ribosomal subunit protein eL43A | Q9HGL8 |
| BN | Large ribosomal subunit protein uL2C | P0CT72 |
| BO | Large ribosomal subunit protein uL3A | P40372 |
| BP | Large ribosomal subunit protein uL4A | P35679 |
| BQ | Large ribosomal subunit protein uL18B | O74306 |

| | | |
|---|---|---|
| BR | Large ribosomal subunit protein eL6 | P79071 |
| BS | Large ribosomal subunit protein uL30C | O60143 |
| BT | Large ribosomal subunit protein eL8 | O13672 |
| BU | Large ribosomal subunit protein uL6B | O74905 |
| BV | Large ribosomal subunit protein uL16A | Q09127 |
| BW | Large ribosomal subunit protein uL5A | P0CT77 |
| BX | Large ribosomal subunit protein uL13 | O74175 |
| BY | Large ribosomal subunit protein eL14 | O94238 |
| BZ | Large ribosomal subunit protein eL15B | Q9US22 |
| Ba | Large ribosomal subunit protein uL13B | O42991 |
| Bb | Large ribosomal subunit protein uL22A | O14339 |
| Bc | Large ribosomal subunit protein eL18B | Q8TFH1 |
| Bd | Large ribosomal subunit protein eL19B | O42699 |
| Be | Large ribosomal subunit protein eL20A | P0CT68 |
| Bf | Large ribosomal subunit protein eL21B | O42706 |
| Bg | Large ribosomal subunit protein eL22 | Q09668 |
| Bh | Large ribosomal subunit protein uL14B | P0CT61 |
| Bi | Large ribosomal subunit protein eL24B | O74884 |
| Bj | Large ribosomal subunit protein uL23A | Q10330 |
| Bk | Large ribosomal subunit protein uL24 | P78946 |
| Bl | Large ribosomal subunit protein eL27A | O14388 |
| Bm | Large ribosomal subunit protein uL15B | P57728 |
| Bn | Large ribosomal subunit protein eL29 | Q92366 |
| Bo | Large ribosomal subunit protein eL30A | P52808 |
| Bp | Large ribosomal subunit protein eL31 | Q9URX6 |
| Bq | Large ribosomal subunit protein eL32A | P79015 |
| Br | Large ribosomal subunit protein eL33B | Q9USG6 |
| Bs | Large ribosomal subunit protein eL34B | Q9URT8 |
| Bt | Large ribosomal subunit protein uL29 | O74904 |
| Bu | Large ribosomal subunit protein eL36B | O94658 |
| Bv | Large ribosomal subunit protein eL37B | P05733 |
| Bw | Large ribosomal subunit protein eL38 | P49167 |
| Bx | Large ribosomal subunit protein eL39 | P05767 |
| By | Ubiquitin-ribosomal protein eL40A fusion protein | P0CH06 |

| **Chain ID** | **Chain name** | **NCBI Reference** |
|---|---|---|
| AA | 18S small subunit ribosomal RNA | NR_151432.1 |
| B2 | 28S large subunit ribosomal RNA | NR_151433.1 |
| B3 | 5S large subunit ribosomal RNA | NR_151410.1 |
| B4 | 5.8S large subunit ribosomal RNA | NR_151436.1 |

**Table S3. Cryo-ET data collection and structure refinement statistics.**

| ***S. pombe*** **strain** | **WT** | **Δ*cpc2*** | **Δ*dnm1*** |
|---|---|---|---|
| **Data collection and processing** | | | |
| Microscope | Titan Krios G3i | Titan Krios G4i | Titan Krios G4i |
| Voltage (kV) | 300 | 300 | 300 |
| Energy filter | Bioquantum | Selectris X | Selectris X |
| Energy slit (eV) | 10 | 10 | 10 |
| Detector | K3 | Falcon 4i | Falcon 4i |
| Detection mode | Counting | Counting | Counting |
| Nominal magnification | 42,000x | 42,000x | 64,000x |
| Pixel size (Å) | 2.075 | 3.033 | 1.871 |
| Defocus range (μm) | -2/-4 | -3/-6 | -3/-5 |
| Defocus step (μm) | 0.5 | 0.5 | 0.5 |
| Tilt acquisition scheme | Dose-symmetric | Dose-symmetric | Dose-symmetric |
| Tilt range (°) | -63/+45 | -43/+59 | -44/+60 |
| Tilt step (°) | 2 | 3 | 2 |
| Tilt exposure ($e^{-}$/Å$^2$/tilt) | 2.2 | 3.0 | 2.4 |
| Total dose ($e^{-}$/Å$^2$) | 121 | 105 | 127 |
| Number of tilts per series | 55 | 35 | 53 |
| Movie format | .tif | .eer | .eer |
| Number of frames per tilt | 9 | 504 | 162 |
| Number of acquired TS | 125 | 12 | 44 |
| Number of processed TS | 53 | 11 | 11 |
| Map resolution at FSC=0.143 (Å) | 11 | 45 | 33 |

**Movie S1. WT *S. pombe* cells are round and decorated with ribosome arrays at day 7 of glucose depletion.**

3D rendering of a segmented cryo-ET tomogram of a mitochondrion in a WT *S. pombe* cell at day 7 of glucose depletion. Segmentation of the mitochondrion shown in Fig. 2, superimposed with the position and orientation of ribosomes aligned by STA. The OMM is depicted in red and the IMM and its cristae are shown in orange. Ribosomal proteins and RNAs of the LSU and SSU are colored cyan and yellow, respectively. In the second half of the movie, ribosome pentamers, open tetramers, closed tetramers and closed trimers light up in magenta, yellow, green and orange respectively.

**Movie S2. Δ*cpc2 S. pombe* cells are round and not decorated with ribosome arrays at day 7 of glucose depletion.**

3D rendering of a segmented cryo-ET tomogram of a mitochondrion in a Δ*cpc2 S. pombe* cell at day 7 of glucose depletion. Segmentation of the mitochondrion shown in Fig. 4, superimposed with the position and orientation of ribosomes aligned by STA. The OMM is depicted in red and the IMM and its cristae are shown in orange. Ribosomal proteins and RNAs of the LSU and SSU are colored cyan and yellow, respectively.

**Movie S3. Δ*dnm1 S. pombe* cells are elongated and decorated with ribosome arrays at day 7 of glucose depletion.**

3D rendering of a segmented cryo-ET tomogram of a mitochondrion in a *Δdnm1 S. pombe* cell at day 7 of glucose depletion. Segmentation of the mitochondrion shown in Fig. 5, superimposed with the position and orientation of ribosomes aligned by STA. The OMM is depicted in red and the IMM and its cristae are shown in orange. Ribosomal proteins and RNAs of the LSU and SSU are colored cyan and yellow, respectively.

**Movie S4-5: PC1 depicting SSU dynamics for the consensus map using cryoDRGN.**

Analysis of the conformational heterogeneity of the SSU in the consensus map of the S. pombe ribosome from *S. pombe* cell grown in glucose depleted EMM. PC1 and PC2 captured the rotation and rolling conformational changes of the SSU, respectively.

**Movie S6: PC1 depicting SSU dynamics for the focused map using cryoDRGN.**

Analysis of the conformational heterogeneity of the SSU in the inactive map of the S. pombe ribosome depicted a conformational change in H69 in the PTC from *S. pombe* cell grown in glucose depleted EMM, depicting the rotation of the SSU.
